# Primate-specific BTN3A2 protects against SARS-CoV-2 infection by interacting with and reducing ACE2

**DOI:** 10.1101/2024.01.13.575537

**Authors:** Ling Xu, Dandan Yu, Min Xu, Yamin Liu, Lu-Xiu Yang, Qing-Cui Zou, Xiao-Li Feng, Ming-Hua Li, Nengyin Sheng, Yong-Gang Yao

## Abstract

**Background:** Coronavirus disease 2019 (COVID-19) is an immune-related disorder caused by severe acute respiratory syndrome coronavirus 2 (SARS-CoV-2). The complete pathogenesis of the virus remains to be determined. Unraveling the molecular mechanisms governing SARS-CoV-2 interactions with host cells is crucial for the formulation of effective prophylactic measures and the advancement of COVID-19 therapeutics.

**Methods:** We analyzed human lung single-cell RNA sequencing dataset to discern the association of butyrophilin subfamily 3 member A2 (*BTN3A2*) expression with COVID-19. The *BTN3A2* gene edited cell lines and transgenic mice were infected by live SARS-CoV-2 in a biosafety level 3 (BSL-3) laboratory. Immunoprecipitation, flow cytometry, biolayer interferometry and competition ELISA assays were performed in *BTN3A2* gene edited cells. We performed quantitative real-time PCR, histological and/or immunohistochemical analyses for tissue samples from mice with or without SARS-CoV-2 infection.

**Findings:** The *BTN3A2* mRNA level was correlated with COVID-19 severity. *BTN3A2* expression was predominantly identified in epithelial cells, elevated in pathological epithelial cells from COVID-19 patients and co-occurred with *ACE2* expression in the same lung cell subtypes. BTN3A2 targeted the early stage of the viral life cycle by inhibiting SARS-CoV-2 attachment through interactions with the receptor-binding domain (RBD) of the Spike protein and ACE2. BTN3A2 inhibited ACE2-mediated SARS-CoV-2 infection by reducing ACE2 *in vitro* and *in vivo*.

**Interpretation:** These results reveal a key role of BTN3A2 in the fight against COVID-19. Identifying potential monoclonal antibodies which mimic BTN3A2 may facilitate disruption of SARS-CoV-2 infection, providing a therapeutic avenue for COVID-19.

**Funding:** This study was supported by the National Natural Science Foundation of China (32070569, U1902215, and 32371017), the CAS “Light of West China” Program, and Yunnan Province (202305AH340006).

**Research in context:** *Evidence before this study:* Our understanding of the pathogenesis of COVID-19, especially key molecular events in the early stage of viral infection, remains incompletely albeit we witnessed many progresses. This knowledge gap hinders the finding for effective and specific antiviral agents against SARS-CoV-2. The entry of SARS-CoV-2 is mediated by the entry receptor angiotensin-converting enzyme 2 (ACE2) and is affected by host antiviral defenses. Targeting these universal host factors required for virus replication is the most promising approach for effective prevention and treatment of COVID-19.

*Added value of this study:* Our study revealed that *BTN3A2,* a primate-specific gene, acts as a potent inhibitor of SARS-CoV-2 infection *in vitro* and *in vivo*. The up-regulation of BTN3A2 upon SARS-CoV-2 infection competed with the ACE2 receptor for binding to the Spike protein, subsequently reducing ACE2 expression and ACE2-mediated SARS-CoV-2 entry.

*Implications of all the available evidence:* These data highlighted that BTN3A2 as a novel host factor with protective effects against SARS-CoV-2 infection. The BTN3A2 holds considerable potential as a therapeutic drug for mitigating the impact of SARS-CoV-2 and its variants of concern (VOCs).

## Introduction

The emergence of coronavirus disease 2019 (COVID-19), caused by severe acute respiratory syndrome coronavirus 2 (SARS-CoV-2), precipitated a global pandemic with persistent pathological sequelae ^1, 2^. According to the World Health Organization, the pandemic resulted in more than 6.9 million deaths in recent years ^3^, posing a significant threat to global health and economic security ^4^. SARS-CoV-2 induces extensive damage to multiple organ systems via an overactive host immune response ^5, 6^. To date, a lot of research efforts have focused on the development of pan-sarbecovirus vaccines and neutralizing antibodies ^7^. However, the pathogenesis of COVID-19, especially key molecular events in the early stage of viral infection, remains incompletely understood. This knowledge gap contributes to the limited availability of effective and specific antiviral agents against SARS-CoV-2 ^8–10^. Therefore, further efforts are required to clarify host-virus interactions and viral pathogenesis and thus identify candidate targets for the prevention and treatment of SARS-CoV-2 infection.

Receptor binding constitutes a critical step in viral invasion ^11, 12^. In the case of SARS-CoV-2, the Spike protein, specifically the receptor-binding domain (RBD) in S1 subunit, mediates recognition of the host ACE2 receptor to facilitate viral entry ^12–17^. Mutations in the Spike protein, particularly in the RBD, can lead to immune evasion, reduce efficacy of existing vaccines, and escape from neutralizing antibodies, as exemplified by the global spread of the Omicron variants ^18–20^. Consequently, identifying broad-spectrum antiviral targets and additional candidates that are effective against different viral variants has become a critical objective in combating the COVID-19 pandemic. Viral host receptors such as ACE2, which is indispensable for SARS-CoV-2 cellular entry, present attractive candidates for intervention ^14, 21, 22^. ACE2 is a transmembrane carboxypeptidase with broad substrate specificity, including angiotensin II, which is essential for viral entry ^15^. Importantly, as ACE2 is a host factor, its expression is less likely to be affected by viral mutations. This suggests that therapeutic approaches aimed at modulating ACE2 expression may be effective against multiple SARS-CoV-2 variants ^23^. Hence, the exploration of drugs or targets that either directly or indirectly affect ACE2 to modulate SARS-CoV-2 infection represents a viable strategy for investigation.

The butyrophilin subfamily 3 (*BTN3*) is a member of the immunoglobulin transmembrane superfamily ^24, 25^, which is gaining increased attention. The three primary members of the BTN3A family, BTN3A1, BTN3A2, and BTN3A3, share similar extracellular domains. BTN3A1 and BTN3A3 contain an intracellular B30.2 domain that is absent in BTN3A2. The *BTN3A2* is a primate-specific gene that likely evolved during primate lineage development ^26^. Members of the BTN3 family members are expressed in most human immune cell subsets and are known to participate in immunomodulatory functions ^27^. Recent studies reiterated that BTN3A1 plays a key role in T cell function ^28–30^. BTN3A2, in particular, has been implicated in the evolutionary and development of the primate brain ^31^ and is associated with various psychological disorders ^26, 32–34^. While fewer literatures demonstrated an association of this family member with antiviral properties, recent studies have identified BTN3A3 as a restriction factor against Ebola virus replicons ^35^ and as a key barrier to avian influenza A virus spillover into humans ^36^, suggesting a potential involvement of the BTN3A family in modulating host antiviral defense. We got to know the *BTN3A2* gene during our initial integrative analyses for the genome-wide association data of schizophrenia ^26^ and major depressive disorder ^34^, and this gene was featured as a risk gene for these mental disorders. We accidently found that overexpression of BTN3A2 could inhibit SARS-CoV-2 replication in Calu-3 cells when we studied SARS-CoV-2 ^37–42^. Reanalysis of reported data in a subsequent study by Li et al. ^43^ showed up-regulation of the mRNA levels of *BTN3A2* and *BTN3A3*, along with other related genes in Calu-3 cells upon SARS-CoV-2 infection. All these observations suggested that BTN3A2 may play a potent role in host antiviral defense against SARS-CoV-2, though the precise mechanism underlying this regulation remains to be clarified.

In this study, we assessed the potential role of BTN3A2 as a key host factor in combating SARS-CoV-2 infection. Based on publicly available datasets and newly obtained data, we observed an up-regulation of *BTN3A2* mRNA expression in COVID-19 patients and non-human primates (NHPs). Using cellular and animal models of SARS-CoV-2 infection, we also established that BTN3A2 competed with the ACE2 via directly interacting with the SARS-CoV-2 Spike RBD, thereby reducing ACE2 receptor expression and ACE2-mediated SARS-CoV-2 entry. This study identified BTN3A2 as a host factor against SARS-CoV-2 infection, providing a basis for the development of potential small molecule inhibitors targeting BTN3A2 for the treatment of COVID-19.

## Materials and Methods

### Ethics statement

The ethics review board of the Kunming Institute of Zoology (KIZ), Chinese Academy of Sciences (CAS) approved this study. All animal experiments were approved by the Ethics Committee of KIZ, CAS (approval number IACUC-RE-2023-03-004). All experiments involving live SARS-CoV-2 were conducted in the animal biosafety level 3 (ABSL-3) facility at KIZ, CAS (approval numbers KIZP3-XMZR-2021-23 and KIZP3-XMZR-2023-04).

### Viral strains and cells

The SARS-CoV-2 prototype was kindly provided by the Guangdong Provincial Center for Disease Control and Prevention (Guangdong, China), as described in our previous studies ^37–39^. The virus was amplified in Vero-E6 cells and median tissue culture infective dose (TCID_50_) was used to assess viral infectivity. The viral sequence is accessible from the China National Microbiology Data Center (NMDC, Accession No. NMDCN0000HUI). The SARS-CoV-2 variant BA.2 was isolated from a patient in Yunnan Provincial Infectious Disease Hospital and its sequence is accessible at NMDC (Accession No. NMDC60046377) ^44^. Virions from the fourth passage were used in all experiments. The VSV-G pseudotyped viruses (G*ΔG-VSV-Rluc), containing prototype and variant BA.1 ^44^, were kindly provided by Prof. Yong-Tang Zheng at KIZ, CAS. The VSV-G pseudotyped viruses (VSV*ΔG-GFP), co-expressing SARS-CoV-2 spike protein and a green fluorescent protein (GFP) reporter ^42^, were kindly provided by Prof. Qihui Wang at Institute of Microbiology, CAS. All viruses were preserved at −80°C freezer.

The HEK293T, Huh7, A549, and Vero E6 cells were obtained from the Kunming Cell Bank, KIZ, and were grown in Dulbecco’s Modified Eagle Medium (DMEM, high glucose;11965-092, Gibco-BRL, USA) supplemented with 10% fetal bovine serum (FBS, Gibco-BRL,10099-141) and 1% penicillin/streptomycin (15140122, Gibco-BRL, USA). The Calu-3 cells were purchased from Procell Life Science & Technology Co., Ltd, and were cultured in Minimum Essential Medium (MEM; 11095080, Gibco-BRL, USA) supplemented with 10% FBS (Gibco-BRL,USA), 1% MEM non-essential acids (11140050, Gibco-BRL, USA), 1% sodium pyruvate (100 mM, 11360070, Gibco-BRL, USA), and 1% antibiotics. All cells were cultured at 37 °C in a humidified atmosphere of 5% CO_2_, and were used for different purposes of testing.

### Plasmid construction and transfection

Two transcripts of human *BTN3A2* (encoding BTN3A2 long isoform [BTN3A2-L] and short isoform [BTN3A2-S]) were cloned into Flag-tagged pCMV-3Tag-8, as described in our previous report ^26^. Briefly, the coding region of *BTN3A2* was sub-cloned into a pCMV-3Tag-8-His vector using the same primers and sub-cloned into pCMV-HA (Clontech, PT3283-5) using *Eco*R I/*Xho* I (New England Biolabs, USA). The expression vector for the SARS-CoV-2 Spike protein (pCMV3-2019-nCoV-Spike (S1+S2)-long) was purchased from Sino Biological (VG40589-UT, China). The expression vector for ACE2 was purchased from Miaolinbio (pEnCMV-ACE2 (human)-3×FLAG, P16698, China). The expression vectors for the Spike S1 (S-S1) and Spike S2 subunits (S-S2) were reported in our previous study ^45^. The expression vector for the prototype RBD (pEnCMV-RBD (SARS-CoV-2)-3×FLAG) was purchased from Miaolinbio (P15640, China), while the expression vectors for the SARS-CoV-2 variant RBDs of the S1 subunit (including α, γ, δ, BA.1, BA.2, BA.2.12, and BA.5) were kindly provided by Prof. Jinghua Yan at Institute of Microbiology, CAS. All primers used to construct plasmids were listed in Supplementary Table S1. Each construct was verified by direct Sanger sequencing.

The HEK293T (ACE2-293T) and A549 (ACE2-A549) cells with stable expression of ACE2 were transfected using Lipofectamine™ 3000 (L3000008, Invitrogen, USA) following the manufacturer’s protocols to test the antiviral activity during SARS-CoV-2 infection. In brief, the cells were cultured in 12-well plates and grown to 70% confluence. The cells in each well were then transfected with 125 μL of solution containing 1 μg of plasmid DNA, 125 μL of Opti-MEM medium (31985-070, Gibco-BRL,USA), and 5 μL of Lipofectamine™ 3000. Cells were incubated with the transfection mixture for 6 h at 37 °C in a humidified atmosphere of 5% CO_2_, then the medium changed to growth medium.

### Generation of genetically modified cell lines

We used Huh7 and Calu-3 cells, which have a moderate expression of *BTN3A2* mRNA and can be naturally infected by SARS-CoV-2, to establish cell lines with BTN3A2 knockout. In brief, two single-guide RNAs (sgRNAs) (sgBTN3A2 #1: TGTTCTCTCCCTTGGCGTTGCTCCACTGTA; sgBTN3A2 #3:ATCATGAGAGGCGGCTCCGGGGAGGGTGTATC) targeting different coding regions of the *BTN3A2* gene were cloned into linearized lentiCRISPRv2 (#52961, Addgene, USA). Lipofectamine™ 3000 transfection reagent in Opti-MEM medium was used to co-transfect HEK293T cells with psPAX2 (#12260, Addgene, USA), pMD2.G (# 12259, Addgene, USA), and indicated individual sgRNA constructs at a ratio of 2:1:3, in accordance with the manufacturer’s instructions. Transfection medium was replaced with viral production medium at 6 h after transfection. The supernatants of the respectively transfected HEK293T cells were collected at 72 h after transfection and filtered using a 0.45 μm filter (SLHV033RB, Millipore, USA). Huh7 and Calu-3 cells were transduced with 1 mL of the above harvested lentiviral stock with polybrene (1 mg/mL, H8761, Solarbio, China) for 24 h. The cells were then switched to 2 mL of fresh medium and puromycin (1 μg/mL, A1113803, Gibco-BRL, USA) to select for successfully transduced Huh7 and Calu-3 cells. We obtained two strains of Huh7 cell with BTN3A2 knockout (sgBTN3A2-1 Huh7 cell and sgBTN3A2-3 Huh7 cells) and two strains of Calu-3 cell with BTN3A2 knockout (sgBTN3A2-1 Calu-3 cell and sgBTN3A2-3 Calu-3 cell), respectively, after verification by using Western blot for the knockout of BTN3A2. The lentiviral particle (sgNC; generated by transfecting HEK293T cells with the lentiCRISPRv2 empty vector, psPAX2, and pMD2.G) was used to establish control cell lines (sgNC Huh7 cells and sgNC Calu-3 cells) for those Huh7 cell and Calu-3 cells with BTN3A2 knockout.

We also established Huh7 cell and Calu-3 cells with stable expression of the BTN3A2 isoforms (BTN3A2-L and BTN3A2-S) ^26^ using the TETon system (632162, Clontech, USA), respectively. Briefly, the coding region sequence of the BTN3A2-L or BTN3A2-S was sub-cloned from pCMV-HA-BTN3A2-L and pCMV-HA-BTN3A2-S expression vectors into the pLVX-tight-puro vector (632162, Clontech, USA) (Supplementary Table S1). The TETon system was combined with a separate lentiviral compatible vector containing the reverse tetracycline-controlled transactivator Advance (rtTA Advance) (632163, Clontech, USA). Lentiviral particles corresponding to either TETon system or rtTA Advance were generated using the above procedures. Huh7 and Calu-3 cells were co-transduced with the TETon system containing the indicated targets (pLVX-Vector to make control cells, pLVX-BTN3A2-L and pLVX-BTN3A2-S for overexpression of BTN3A2-L and BTN3A2-S, respectively) and rtTA Advance lentiviral particles (each 1 mL) in the presence of polybrene (1 mg/mL) for 48 h. The culture medium was replaced with 2 mL of fresh medium and incubated for an additional 24 h. Cells were subsequently selected using G418 (250 ng/mL, A1720, Sigma-Aldrich) and puromycin (1 μg/mL) for a minimum of two weeks. We obtained a total of four strains with stable expression of BTN3A2 isoform (Huh7 cells: BTN3A2-L Huh7 and BTN3A2-S Huh7; Calu-3 cells: BTN3A2-L Calu-3 and BTN3A2-S Calu-3) and two strains of control cells (Vector Huh7 and Vector Calu-3).

To generate cell lines with a stable ACE2 expression, we used HEK293T cells and A549 cells, as both have a high transfection efficiency and a high SARS-CoV-2 infection efficiency after stable overexpression of ACE2. Briefly, a lentiviral expression system for ACE2 was established using the pLenti-CMV-ACE2-Flag plasmid (Public Protein/Plasmid Library, China) in HEK293T cells, and viral particles in the supernatants were collected. For lentiviral expression of ACE2, both HEK293T and A549 cells were seeded in 6-well plates and transduced by infection mixture containing 1 mL of culture medium with 1 μg of polybrene (H8761, Solarbio, China) and 1 mL of viral supernatants. Subsequently, the culture medium was replaced with 2 mL of fresh medium containing puromycin (1 μg/mL) to select for successfully transduced HEK293T and A549 cells overexpressing ACE2. The cell lines with stable expression of ACE2 (ACE2-A549 cells and ACE2-HEK293T cells) were verified by Western blot.

### Western blotting and immunoprecipitation

Western blotting and immunoprecipitation were performed using the same procedures described in our previous studies ^46, 47^. In brief, cells were lysed on ice in RIPA lysis buffer (P0013, Beyotime Institute of Biotechnology, China) and centrifuged at 12 000 *g* for 5 min at 4 °C to harvest cell lysates. Protein concentration was determined using a BCA Protein Assay Kit (P0012, Beyotime Institute of Biotechnology, China). Bovine serum albumin (BSA; P0007, Beyotime Institute of Biotechnology, China) was used as the protein standard. Cell lysates (20 μg of total protein/sample) were separated by 12% sodium dodecyl-sulfate polyacrylamide gel electrophoresis (SDS-PAGE), before transfer to polyvinylidene fluoride (PVDF) membranes (IPVH00010, Roche Diagnostics, China) using standard procedures. The membranes were then blocked with 5% non-fat dry milk in Tris-buffered saline (#9997, Cell Signaling Technology, USA) with 0.1% Tween 20 (P1379, Sigma, USA) (TBST) at room temperature for 2 h. Membranes were incubated with primary antibodies against His (1:5 000, M20001M, Abmart, China), HA (1:5 000, M20003M, Abmart, China), Flag (1:5 000, M20008M, Abmart, China), ACE2 (1:1 000, 66699-1-Ig, Proteintech, China), BTN3A2 (1:1 000, TA500730, Origene, USA), Spike (1:1 000, 28869-1-AP, Proteintech, China), and Tubulin (1:10 000, OTI3H2, ZSGB-Bio, China) overnight at 4 °C, respectively. The membranes were washed three times (5 min each) with TBST and incubated with TBST-conjugated anti-mouse (5450-0011) or anti-rabbit (5220-0458) secondary antibody (1:10 000, Milford, MA, USA) for 1 h at room temperature. After washing with TBST three times, the proteins on the membranes were detected by enhanced chemiluminescence reagents (WBKLS0500, Millipore, USA). All antibodies have been validated in our previous studies ^37, 38, 46, 48^ or using the western blot assays in this study.

For immunoprecipitation, appropriate antibodies were incubated with protein G-agarose beads (15920010, Life Technologies, USA) to form a complex for 2 h at room temperature. Cells were lysed with RIPA lysis buffer on ice for 1 h, followed by centrifugation at 12 000 *g* for 10 min at 4 °C. The lysates were then immunoprecipitated with the complex at 4 °C overnight, followed by four washes with RIPA lysis buffer and resuspension in loading sample buffer for SDS-PAGE.

### Syncytia formation assay

The syncytia formation assay was performed by using a co-culture system of Huh7 cells and HEK293T cells, following a previously described procedure ^49, 50^. Briefly, Huh7 cells with overexpression of BTN3A2-L or BTN3A2-S, and control cells (Vector Huh7 cells) were seeded in a 4-well chamber slide (Nunc™ Lab-Tek™ II, 177399, Thermo Scientific, USA) at a density of 1×10^4^ cells/ well for overnight culture, then were transfected with the pX-330 plasmid (Addgene) (500 ng/well) using Lipofectamine™ 3000 (L3000008, Invitrogen, USA) for 16 h to be ready for the syncytia formation assay. The HEK293T cells were cultured in a 12-well plate to approach 60-70% confluence and were transfected with expression vector for Spike of SARS-CoV-2 Delta ^44^ (1 μg) for 16 h, then cells were trypsinized and re-suspended in DMEM supplemented with 5% FBS at a density of 2×10^4^ cells/mL. We seeded HEK293 cell suspension (500 µL/well) into the 4-well chamber slide with the above prepared Huh7 cells (BTN3A2-L, BTN3A2-S, and Vector Huh7 cells) for co-culture for 20 h. Cells were collected and fixed with 4% paraformaldehyde (PFA, 158127, Sigma-Aldrich, USA) for further immunofluorescence assay.

### Immunofluorescence

Huh7 cells with BTN3A2 overexpression (BTN3A2-L and BTN3A2-S Huh7 cells) or knockout (sgBTN3A2-1 and sgBTN3A2-3 Huh7 cells) and corresponding control cells (Vector Huh7 and sgNC Huh7) were seeded on glass coverslips and grown overnight to 40% confluence in DMEM supplemented with 10% FBS at 37 °C in 5% CO_2_. The BTN3A2-L and BTN3A2-S Huh7 cells were treated with Dox (1 mg/mL, D9891, Sigma-Aldrich, USA) for 24 h to induce the expression of BTN3A2 isoform. All cultured cells were infected with the VSV*ΔG-GFP SARS-CoV-2 S for 24 h before harvest for immunofluorescence analysis. Cells were fixed with 4% paraformaldehyde for 10 min and were incubated with mouse anti-BTN3A2 (1:50) overnight at 4 °C. After three washes with phosphate-buffered saline (PBS) with 0.1% Tween 20 (PBST; 5 min each), immunoreactivity was detected by using the FITC-conjugated secondary antibody (1:500; KPL, 172-1506; incubation for 1 h at room temperature). Nuclei were counterstained with 1 μg/mL DAPI (Roche Diagnostics, 10236276001) and the slides were visualized under an Olympus FluoView™ 1000 confocal microscope.

### Flow cytometry

To measure the cell surface expression level of ACE2, Huh7 cells and Calu-3 cells with BTN3A2 overexpression (BTN3A2-L Huh7 cells, BTN3A2-S Huh7 cells, BTN3A2-L Calu-3 cells and BTN3A2-S Calu-3 cells) or knockout (sgBTN3A2-1 Huh7 cells, sgBTN3A2-3 Huh7 cells, sgBTN3A2-1 Calu-3 cells, and sgBTN3A2-3 Calu-3 cells), and corresponding control cells (Vector Huh7, Vector Calu-3, sgNC Huh7, and sgNC Calu-3) were washed with PBS and stained with mouse anti-ACE2 (66699-1-Ig, Proteintech, China) at a 1:50 dilution for 30 min at 4 °C. The cells were then washed twice and re-suspended in 1×PBS containing secondary antibody (AF647-labeled donkey anti-goat immunoglobulin G (IgG), A31571, Invitrogen, USA) at a 1:500 dilution. After 30 min of incubation at 4 °C, the cells were washed twice and re-suspended in 1×PBS before flow cytometry analysis using FACSCelesta (BD Biosciences, USA). Data were analyzed using FlowJo software.

### Re-analysis of RNA-sequencing (RNA-seq) datasets

RNA-seq data of blood leukocytes collected from 26 hospitalized non-COVID-19 patients and 100 COVID-19 patients with different degrees of disease severity were obtained from the Gene Expression OmniBus (GEO, www.ncbi.nlm.nih.gov/geo/, accession number GSE157103) ^51^. Of these individuals, 16 non-COVID-19 patients and 50 COVID-19 patients were admitted to the intensive care unit (ICU). The RNA-seq dataset of monocytic-myeloid-derived suppressor cells (M-MDSCs) from 12 COVID-19 patients (four severe, four asymptomatic, and four convalescent) and four controls were downloaded from the GEO database (accession number GSE178824) ^52^. Raw RNA-seq data of 15 tissues from three rhesus macaques (*Macaca mulatta*) infected with SARS-CoV-2 and one control were downloaded from the China National Center for Bioinformation (https://www.cncb.ac.cn/) (accession number CRA004025) ^53^. The raw sequencing reads were first cleaned using Trimmomatic v0.33 ^54^, then mapped to the rhesus macaque reference genome (Mmul_10, https://www.ncbi.nlm.nih.gov/datasets/genome/GCF_003339765.1/) using STAR v2.7.3a ^55^. Gene expression quantification was conducted with RSEM v1.3.1 ^56^.

Single-cell RNA-seq (scRNA-seq) data of peripheral blood from five healthy donors and 27 COVID-19 patients were downloaded from https://atlas.fredhutch.org/fredhutch/covid/dataset/merged ^57^. ScRNA-seq data of lung tissues from seven controls and 19 COVID-19 patients were downloaded from https://singlecell.broadinstitute.org/single_cell/study/SCP1219/columbia-university-nyp-covid-19-lung-atlas ^58^. The expression levels of *BTN3A2* and *ACE2* in bulk non-disease human tissues and single cells were obtained from the GTEx Portal (https://www.gtexportal.org/) ^59, 60^. RNA-seq data of mock (*n*=3) and SARS-CoV-2 (*n*=3)-infected lung-derived Calu-3 cells, primary human lung epithelial cells (NHBE) and lung biopsies from healthy controls (*n*=2) and postmortem COVID-19 patients (*n*=2) were obtained from the GEO database (accession number GSE147507) ^61, 62^. Raw counts were normalized using DESeq2 ^63^ and batch effects among samples were removed using the removeBatchEffect function from limma ^64^. For bulk RNA-seq data, the *BTN3A2* expression levels were compared between different groups using two-sided *t*-tests. For scRNA-seq data, the *BTN3A2* expression levels were compared between different groups for each cell type using two-sided Wilcox rank sum tests. Expressional correlations between two genes were calculated using Pearson’s correlation.

### Animals and tissues

Left lobe lung tissue from rhesus macaques (*n*=6) ^65^ and cynomolgus macaques (*Macaca fascicularis*; *n*=8) ^66^ infected with SARS-CoV-2 were obtained from the tissue bank of the ABSL-3 facility at KIZ, CAS. Lung tissues from healthy rhesus macaques (*n*=5) and cynomolgus macaques (*n*=6) were purchased from the National Resource Center for Non-Human Primates, KIZ.

C57BL/6J wild-type (WT) mice were obtained from the experimental animal core facility of KIZ, CAS. Humanized *BTN3A2* knock-in mice were generated using CRISPR/Cas9 on a C57BL6/J background by inserting the coding sequence of the human *BTN3A2* gene (Transcript ID: ENST00000356386.6) with a HA-tag in its N-terminus into the *Hipp11* (*H11*) locus. The BTN3A2 transgenic (BTN3A2-tg) mice were created by GemPharmatech (Nanjing, China). Both male and female mice (4 weeks) were used for the experiments. All mice were genotyped by PCR with genomic DNA prepared from tail tips using primer pair 5’-CAGCAAAACCTGGCTGTGGATC-3’ / 5’-CGTAGATGTACTGCCAAGTAGG-3’. Mice were housed in clear plastic cages and bred in a specific pathogen-free condition at the experimental animal core facility of KIZ, with free access to water and food under 22 ± 2 °C, 50 % humidity, and 12 h light/dark cycle.

### Animal models for SARS-CoV-2 infection

To evaluate whether adeno-associated virus (AAV) exhibits good efficiency for overexpressing exogenous protein in mouse tissues, we infected four age-matched C57BL/6J (*n*=2) and BTN3A2-tg (*n*=2) mice with enhanced green fluorescent protein (EGFP)-expressing AAV (AAV-EGFP) (PackG410ene Biotech, Guangzhou, China). Briefly, mice were intranasally infected with AAV-EGFP (PackGene Biotech, Guangzhou, China) at a total volume of 30 μL containing 2×10^12^ viral genome copies. The AAV-EGFP-infected mice were sacrificed at 14 days post-infection (dpi) with AAV.

A total of 6-10 mice per group were used for infection according to our previous study ^41^. Human ACE2 (hACE2) was overexpressed using hACE2-expressing AAV (AAV-hACE2) in C57BL/6J (*n*=9) mice and BTN3A2-tg mice (*n*=8). In brief, mice were intranasally infected with AAV-hACE2 (PackGene Biotech, Guangzhou, China) at a total volume of 30 μL containing 2×10^12^ genome copies. The C57BL/6J (*n*=7) and BTN3A2-tg (*n*=6) mice were monitored daily until SARS-CoV-2 infection at 14 dpi with AAV-hACE2. After anesthetization with isoflurane (RWD Life Science, China), the mice were intranasally infected with a total volume of 30 μL containing 1×10^5^ TCID_50_ of SARS-CoV-2 variant BA.2. All infected animals were housed at the ABSL-3 facility on a 12 h light/dark cycle, with free access to food and water, and were sacrificed at 3 dpi. The remaining C57BL/6J (*n*=2) and BTN3A2-tg (*n*=2) mice with AAV-hACE2 infection but without SARS-CoV-2 infection were assigned as the control group and sacrificed at 14 dpi with AAV.

Tissue samples were collected from all animals after euthanasia and stored in a −80 °C freezer (for quantification of mRNA and protein levels) or in 4% PFA (for histological analysis and immunochemistry) until use.

### Measurement of viral RNAs

RNA copies per mL (cell RNA) or per μg (lung samples) were determined by quantitative real-time polymerase chain reaction (qRT-PCR), as described in our previous study ^38^. Briefly, total RNA was extracted from 200 µL of indicated cell supernatants using a High Pure Viral RNA Kit (11858882001, Roche, Germany). The extracted RNA was eluted in 50 µL of elution buffer. TRIzol Reagent (15596026CN, Thermo Fisher Scientific, USA) was used for total RNA isolation from cells and homogenized tissues. Viral RNA copies were detected using a THUNDERBIRD Probe One-Step qRT-PCR Kit (QRZ-101, TOYOBO, Japan). A total of 25 µL reaction solution, containing 8.7 µL of extracted RNA from indicated cell supernatant or 1 µg of total RNA from cells or lung tissue samples, was prepared for qRT-PCR analysis ^38^. The Applied Biosystems 7500 Real-Time PCR system was used for qRT-PCR analysis with the following thermal cycle conditions: 10 min at 50 °C for reverse transcription, 60 s at 95 °C, followed by 40 cycles at 95 °C for 15 s and 60 °C for 45 s. In each run, serial dilutions of the SARS-CoV-2 RNA reference standard (National Institute of Metrology, China) were used in parallel to calculate copy numbers in each sample.

### Pathological analysis

The lungs of SARS-CoV-2-infected mice were fixed in 4% PFA for 7 days, processed in paraffin (Leica EG1160, Germany), sectioned to 3-4 μm thickness (Leica RM2255, Germany), and stained with hematoxylin and eosin (H&E), as described in our previous studies ^37, 38^. The slices were imaged using a Nikon Eclipse E100 microscope (Japan) and were evaluated in a blinded manner by two pathologists.

Immunohistochemical analysis of lung tissue was performed based on previously described procedures ^37, 48^. In brief, each section was baked at 65 °C for 30 min, followed by deparaffinization using xylene and subsequent hydration with decreasing concentrations of ethanol (100% to 75%). Heat-induced antigen retrieval was performed using sodium citrate buffer (pH 6.0). Endogenous peroxidase was blocked with 3% hydrogen peroxide for 25 min, and non-specific binding was blocked with 3% BSA for 30 min at room temperature. Specimens were incubated overnight at 4 °C with rabbit anti-SARS-CoV-2 nucleocapsid protein (1:200, 26369, Cell Signaling Technology, USA), mouse anti-ACE2 (1:1000, Proteintech,66699-1-Ig), mouse anti-GFP (1:2000, sc-9996, Santa Cruz Biotechnology, USA), and mouse anti-BTN3A2 antibodies (1:50, CF500730, Origene, USA), respectively. The slices were incubated with mouse IgG1 kappa isotype control (1:1000, 14-4714-82, Thermo Fisher Scientific, USA) as a negative control for antigen staining. After washing with PBS with 0.1% Tween 20 three times, the lung slices were incubated with anti-rabbit IgG (GB111738, Servicebio, China) or anti-mouse IgG secondary antibody (GB111739, Servicebio, China) for 50 min at room temperature and visualized with 3,3’-diaminobenzidine tetrahydrochloride. The slices were counterstained with hematoxylin, dehydrated and mounted on a slide, and imaged under an Olympus OlyVIA microscope (Japan).

For immunofluorescence assay, lung tissue sections were blocked in PBST supplemented with 5% BSA for 1 h, followed by an incubation with primary anti-mouse ACE2 (1:1000; Proteintech,66699-1-Ig) and anti-rabbit HA-Tag (1:800; Cell signaling technology, C29F4) for 16 h. After three washes with PBST (each 5 min), sections were incubated with a FITC-conjugated anti-Rabbit IgG (1:500; Life Technologies, A21207) or anti-mouse IgG (1:500; Invitrogen, A21202) secondary antibody, and nuclei were counterstained with DAPI. The slides were visualized under an Olympus FluoView™ 1000 confocal microscope (Olympus, America).

### Enzyme-linked immunosorbent assay (ELISA) competition assays

Spike S1-Fc (6.25 ng/mL, 40591-V02h, Sino Biological, China) was mixed with ACE2-Fc (50 ng/mL, 10108-H02H, Sino Biological, China) or Human IgG-Fc (50 ng/mL, 10702-HNAH, Sino Biological, China) at 4 °C overnight. The mixture was then added with a series of concentrations (0, 6.25, 12.5, 25, 50 ng/mL) of BTN3A2-L-His (13515-H08H, Sino Biological, China). These sample series were transferred to a plate pre-coated with an antibody specific to SARS-CoV-2 S1RBD (E-EL-E605, Elabscience, China). After 2 h of incubation, the plate was washed by wash buffer three times. Avidin-horseradish peroxidase (HRP) conjugate was successively added to each microplate well and incubated at 4 °C for 30 min. The free components were then removed by washing and substrate solution was added to each well. The enzyme-substrate reaction was terminated by the addition of stop solution according to the manufacturer’s instructions. Those wells containing different residual concentration of S1-RBD, biotinylated detection antibody, and Avidin-HRP conjugate appeared blue. Optical density was measured using a BioTek Synergy HT microplate reader (BioTek, USA).

### Biolayer interferometry assay

The biolayer interferometry assays were performed using the Octet RED96 system (ForteBio, Germany). Briefly, ACE2-Fc protein (10 μg/mL) was captured on AR2G Biosensors (18-5095, ForteBio, Germany) and incubated with a series of concentrations of BTN3A2-L-His (62.5, 250, 1 000, and 2 000 nM). The biolayer interferometry assays were performed via four steps: (i) ACE2-Fc protein loading onto AR2G Biosensors (360s); (ii) baseline; (iii) association of BTN3A2-L-His to determine Kon (association rate constant) (360 s); and (iv) dissociation of BTN3A2-L-His to determine Kdis (dissociation constant) (360 s). The baseline and dissociation steps were performed in SD buffer (pH 7.4 PBS, 0.05% Tween 20, 0.01% BSA) and biosensor drifting was corrected by background subtraction. Background wavelength shifts were measured in the reference biosensors only loaded with ACE2-Fc in SD buffer. All steps were conducted in a shaker (1 000 rpm, 30 °C). The data were analyzed and fitted in a 1:1 binding model using Octet data analysis software (ForteBio v9.0). The same procedure was used to determine whether BTN3A2 binds to Spike S1. Briefly, BTN3A2-L-His protein (10 μg/mL) was captured on AR2G Biosensors and incubated with Spike S1-Fc (125, 250, 500, and 1 000 nM) at various concentrations.

We also tested whether BTN3A2-S, BTN3A1 and BTN3A3 bind to Spike S1 or ACE2, respectively, following the above procedure. Briefly, BTN3A2-S-His protein (10 μg/mL, Zoonbio Biotechnology, China) was captured on AR2G Biosensors and incubated with Spike S1-Fc (250 nM, 500 nM, and 1 000 nM) or ACE2-Fc (500 nM, and 1 000 nM) at various concentrations. For biolayer interferometry assays of BTN3A1-His (10 μg/mL, 15973-H08H, Sino Biological, China) and BTN3A3-His (10 μg/mL, 13432-H08H, Sino Biological, China) with Spike S1-Fc (1 000 nM) or with ACE2-Fc (1 000 nM), we only used one concentration because of the limited amount of each protein.

### Statistical analysis

All statistical analyses were performed using GraphPad Prism version 8.0.1 (GraphPad Software, Inc., La Jolla, CA, USA). Comparisons between different groups were conducted by using one-way analysis of variance (ANOVA) with Dunnett’s multiple comparisons. Data are presented as mean ± SD. A *P* value <0.05 was considered as statistically significant.

## Results

### *BTN3A2* was altered in COVID-19 patients and SARS-CoV-2-infected NHPs

We first analyzed *BTN3A2* mRNA expression in peripheral blood mononuclear cells (PBMCs) from COVID-19 patients based on published transcriptomic data ^51^ to examine potential relevance between *BTN3A2* and SARS-CoV-2 infection. Results showed a significantly increased *BTN3A2* mRNA level in COVID-19 patients (Non-ICU COVID-19) compared to non-COVID-19 patients (ICU and Non-ICU) (*P*=5.3×10^-4^, Fig. 1a). However, in critically ill COVID-19 patients (ICU COVID-19), *BTN3A2* mRNA expression was significantly reduced relative to Non-ICU COVID-19 patients (*P*=1.42×10^-6^, Fig. 1a). In M-MDSCs from COVID-19 patients with different disease severities and stages ^52^, no marked differences in *BTN3A2* mRNA expression were found between the healthy controls and asymptomatic patients or between healthy controls and convalescent patients, but expression was significantly reduced in patients with severe COVID-19 relative to healthy controls (*P*=3.6×10^-3^, Fig. 1b). Re-analyses of scRNA-seq data of PBMCs, B cells, T cells (CD4^+^ T, CD8^+^ T, and other T cells), and natural killer (NK) cells ^67^ also showed an up-regulation of *BTN3A2* mRNA expression in most cell types in COVID-19 patients compared to healthy controls (Fig. 1c).

**Fig. 1.**
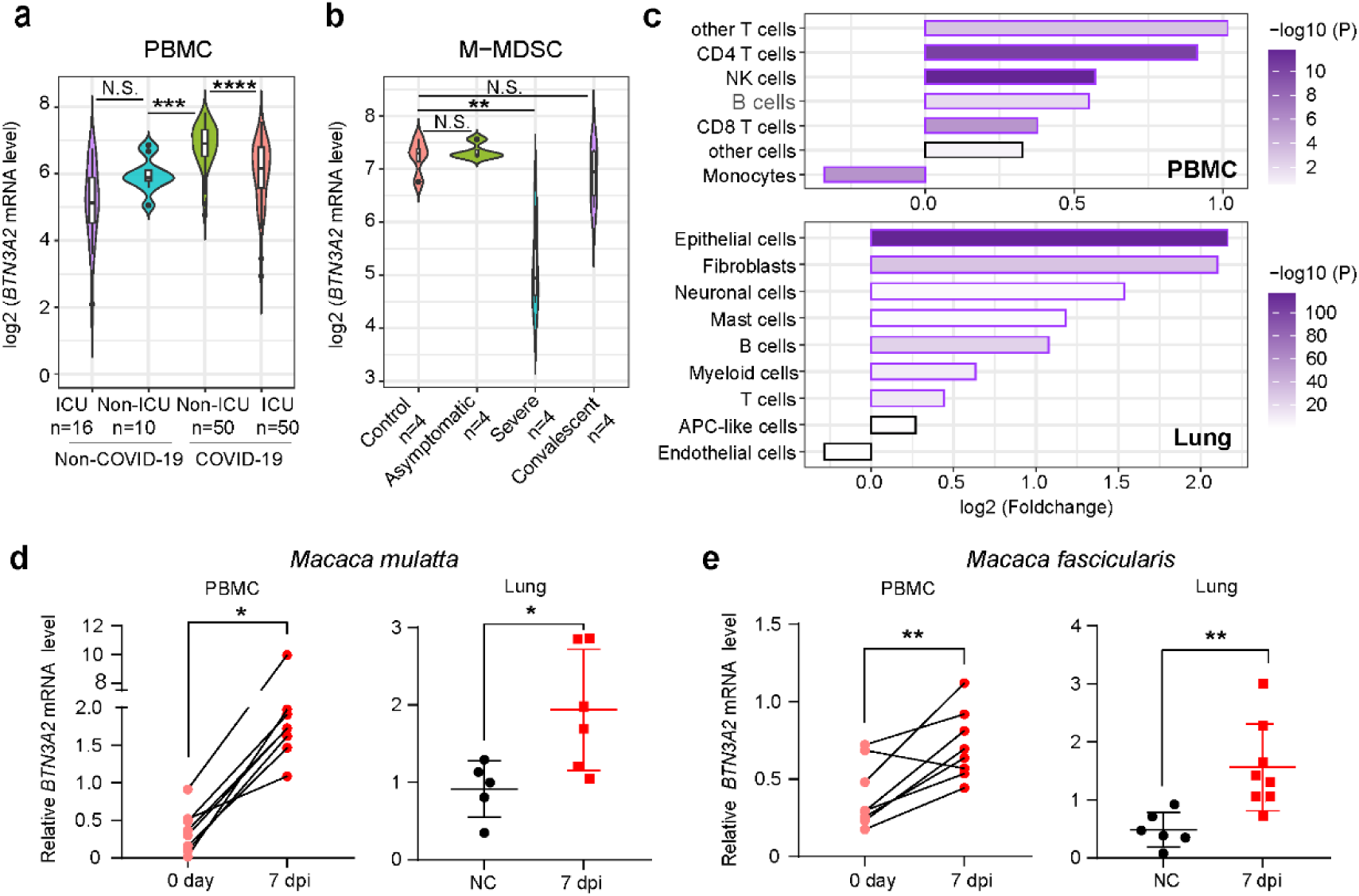
BTN3A2 was up-regulated in COVID-19 patients and NHPs infected with SARS-CoV-2. **a**: Violin plot of *BTN3A2* mRNA levels in peripheral blood mononuclear cells (PBMCs) from COVID-19 patients with (ICU COVID-19) or without ICU experience (Non-ICU COVID-19) compared to healthy individuals (Non-ICU Non-COVID-19) and individuals in ICU but not for COVID-19 (ICU Non-COVID-19). Original dataset (GSE157103) was reported in Overmyer et al. ^51^. **b**: Violin plot of *BTN3A2* mRNA levels in monocytic-myeloid-derived suppressor cells (M-MDSCs) based on the GSE178824 dataset ^52^. *BTN3A2* mRNA level was normalized by trimmed mean of M-values. **a-b**: *BTN3A2* mRNA levels were normalized by transcripts per million (TPM). Gene expression level was determined as log2 fold-change. **c**: Histogram showing up-regulation of *BTN3A2* mRNA levels in PBMCs (*upper*) and lung tissues (*lower*) of COVID-19 patients based on datasets reported in Tian et al. ^57^ and Melms et al. ^58^. Statistical significance was assessed by two-sided Wilcox rank sum test. Purple borders indicate *P*<0.05 and black borders indicate *P*>0.05. Statistical significance was assessed by two-sided *t*-test. **d-e**: *BTN3A2* mRNA levels were up-regulated in PBMCs (*left*) and lung tissues (*right*) of rhesus (**d**) and cynomolgus monkeys (**e**) challenged with or without SARS-CoV-2. For PBMCs, samples were collected before SARS-CoV-2 infection (0 day) and at 7 days post infection (dpi). For lung tissues, samples were collected from animals without SARS-CoV-2 infection (NC) and animals infected with SARS-CoV-2 at 7 dpi. *BTN3A2* mRNA level was measured by qRT-PCR and normalized to *GAPDH*. Differences between two groups were determined by two-tailed Student’s paired *t* test (PBMCs) and unpaired *t* test (Lung), respectively. *, *P* < 0.05; **, *P* < 0.01.

The *BTN3A2* gene is expressed in various myeloid and immune cells and plays an important role in the immune response^24, 27, 68^. However, the functional implications of elevated *BTN3A2* expression in the lungs, which are the primary target organs for SARS-CoV-2, remain to be elucidated. Hence, we first explored the expression of *BTN3A2* in healthy human lung tissues using available GTEx data ^59^. Results indicated that the lungs had the second highest *BTN3A2* expression among 30 tested tissues (Supplementary Fig. S1a). Among the different cell types in the lung tissue, alveolar type 1 and 2 cells, together with alveolar macrophages, showed relatively high *BTN3A2* expression (Supplementary Fig. S1b). Further analysis of the scRNA-seq datasets of COVID-19 patient lungs ^58^ showed that epithelial cells had the highest *BTN3A2* expression (Fig. 1c), which are the primary target cells of SARS-CoV-2 ^69^.

We further explored *BTN3A2* expression patterns in NHPs challenged with SARS-CoV-2, given their heightened resemblance to humans in the cellular identities and frequencies of putative viral target cells ^13, 70, 71^. Among 15 tissues tested in rhesus monkeys ^53^, lung tissues showed the highest expression level of *BTN3A2* (Supplementary Fig. S1c). Immunohistochemical analysis also showed that BTN3A2 was constitutively expressed in the healthy lung tissues of rhesus monkeys (Supplementary Fig. S1d). We quantified the changes in *BTN3A2* mRNA expression in rhesus and cynomolgus monkeys using qRT-PCR. Results indicated that *BTN3A2* mRNA levels were significantly up-regulated in PBMCs and lung tissues at 7 dpi with SARS-CoV-2 in rhesus macaques (Fig. 1d) and cynomolgus monkeys (Fig. 1e). Similar *BTN3A2* mRNA expression patterns have also been observed in the transcriptomic data of rhesus monkeys with early SARS-CoV-2 infection ^53^ (Supplementary Fig. S1e). Collectively, these results indicated that *BTN3A2* is up-regulated in COVID-19 patients and NHPs infected with SARS-CoV-2. In order to determine whether the increased *BTN3A2* mRNA was caused by the induction of the interferon system upon SARS-CoV-2 ^5, 6^, we treated Calu-3 cells with IFN-α, and observed the up-regulation of full-length *BTN3A2* (*BTN3A2-L*) in cells treated with IFN-α (Supplementary Fig. S1f).

### BTN3A2 acted as a potent SARS-CoV-2 restriction factor

BTN3A2, a member of the immunoglobulin superfamily, is implicated in diverse cellular processes ^24^. BTN3A2 contains both full-length BTN3A2 (BTN3A2-L) and truncated BTN3A2 (BTN3A2-S), as identified in our previous report ^26^. To evaluate the potential role of BTN3A2 in inhibiting SARS-CoV-2, we first ectopically expressed the BTN3A family members (BTN3A1, BTN3A3, and BTN3A2) in ACE2-A549 cells (Supplementary Fig. S2a), which were then infected with SARS-CoV-2. Both BTN3A2 isoforms significantly inhibited SARS-CoV-2 viral copy numbers, whereas BTN3A1 and BTN3A3 had no apparent inhibitory effect (Fig. 2a).

**Fig. 2.**
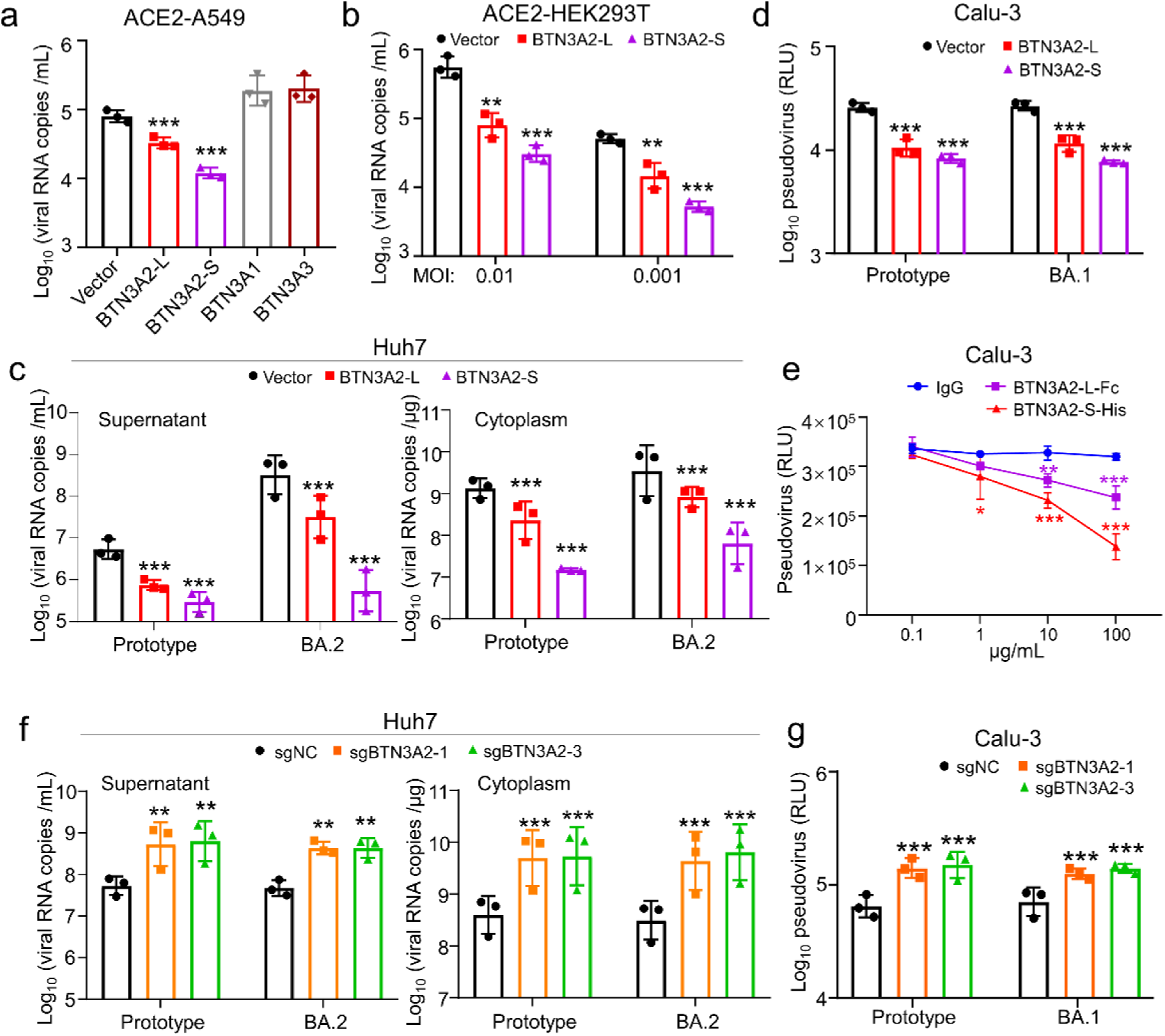
BTN3A2 was identified as a potent restriction factor for SARS-CoV-2 infection. **a**: Overexpression of BTN3A family members inhibited SARS-CoV-2 replication in A549 cells overexpressing human ACE2 (ACE2-A549). ACE2-A549 cells (1×10^5^) were transfected with indicated expression vector (BTN3A2-L, BTN3A2-S, BTN3A1, or BTN3A3) or empty vector (Vector) (each 1 μg) for 24 h, then infected with SARS-CoV-2 (MOI=0.01) for 24 h. Cells were harvested to determine SARS-CoV-2 copies by qRT-PCR. **b**: Overexpression of BTN3A2 inhibited SARS-CoV-2 replication in HEK293T cells overexpressing human ACE2 (ACE2-HEK293T). ACE2-HEK293T cells (5×10^4^) were transfected with indicated expression vector (BTN3A2-L or BTN3A2-S) or empty vector (each 1 μg) for 24 h, then infected with SARS-CoV-2 (MOI=0.01 or 0.001) for 24 h and harvested for quantification of SARS-CoV-2 copies. **c**: Stable expression of BTN3A2 in Huh7 cells inhibited SARS-CoV-2 replication. Huh7 cells overexpressing BTN3A2-L or BTN3A2-S, and control cells (Vector Huh7) (1×10^5^ cells) were treated with DOX (1 μg/mL) for 24 h, then challenged with the SARS-CoV-2 prototype or BA.2 (MOI=1) for 24 h. The culture medium (supernatant, *left*) and cell lysates (cytoplasm, *right*) were collected to quantify SARS-CoV-2 copies. **d**: Stable expression of BTN3A2 in Calu-3 cells inhibited replication of VSV pseudotyped SARS-CoV-2 (prototype and BA.1). The procedure was similar to that in (**c**). After 24h infection, cells were collected, and viral infectivity was quantified based on *Renilla* luciferase activity (RLU) using a *Renilla* Luciferase Assay Kit (Promega, Madison, WI, USA). **e**: Incubation with BTN3A2 isoforms (BTN3A2-L-Fc and BTN3A2-S-His) inhibited replication of VSV pseudotyped SARS-CoV-2 (BA.1) in Calu-3 cells. IgG treatment was used as a negative control. Cells were incubated with BTN3A2 isoform or IgG for 1 h, then infected with VSV pseudotyped SARS-CoV-2 (BA.1) for 24 h and harvested to determine viral infectivity. **f**: Knockout of BTN3A2 increased SARS-CoV-2 replication in Huh7 cells. Huh7 cells with BTN3A2 knockout (sgBTN3A2-1 and sgBTN3A2-3 Huh7) and control cells (sgNC Huh7) were infected with SARS-CoV-2 prototype or BA.2 (MOI=1) for 24 h, then harvested to quantify SARS-CoV-2 copies. **g**: Knockout of BTN3A2 increased VSV pseudotyped SARS-CoV-2 (prototype and BA.1) replication in Calu-3 cells. The procedure for determining viral infectivity was the same as that in (**d**). Data shown in (**b-d**) were average quantitative data from three independent experiments and were presented as mean±standard deviation (SD). Significance was determined by comparing to Vector in each group (a, b, c, and d), IgG control at each dilution (e), or sgNC in each group (f and g). *, *P* <0.05; **, *P*<0.01; ***, *P*<0.001; one-way analysis of variance (ANOVA) with Dunnett’s multiple comparisons.

Compared to a higher multiplicity of infection (MOI), overexpression of BTN3A2 in ACE2-HEK293T cells had a more potent inhibitory effect on lower infectious titers (Fig. 2b and Supplementary Fig. S2b). The Huh7 cell line, which endogenously expresses ACE2 and is naturally susceptible to SARS-CoV-2 ^72^, was used to validate the above results. Overexpressing BTN3A2 in Huh7 cells (Supplementary Fig. S2c) significantly reduced viral copies of the SARS-CoV-2 prototype and BA.2 variant in cytoplasm and culture supernatant (Fig. 2c). Similar inhibitory effects were observed using pseudotyped virus (which is a heterologous virus with Spike protein) ^44^ in BTN3A2-L and BTN3A2-S Calu-3 cells (Fig. 2d). Interestingly, pre-incubation of soluble BTN3A2-L and BTN3A2-S proteins in Calu-3 cells blocked viral entry into cells, with a dose-dependent inhibitory effect (Fig. 2e).

To further test the potential role of endogenous BTN3A2 in controlling SARS-CoV-2 infection, we depleted BTN3A2 expression in Huh7 cells using CRISPR-Cas9 gene editing, as described in our previous study ^47^, with two individual sgRNAs (sgBTN3A2-1 and sgBTN3A2-3; Supplementary Fig. S2d). sgBTN3A2-1 and sgBTN3A2-3 cells were more susceptible to authentic and pseudovirus SARS-CoV-2 than control sgNC cells (Fig. 2f-g). Thus, these findings suggest that BTN3A2 can restrict SARS-CoV-2 infection.

### BTN3A2 directly interacted with Spike protein at RBD

Next, we characterized the effects of BTN3A2 on specific steps of the viral replication cycle. Initially, we assessed whether BTN3A2 inhibits SARS-CoV-2 attachment to the cell surface. Our data showed that SARS-CoV-2 viral copies were significantly reduced in Huh7 cells with stable expression of BTN3A2-L and BTN3A2-S, contrasting to corresponding control cells (Vector Huh7) (Fig. 3a and Supplementary Fig. S3a). This result was confirmed by using a VSV-G pseudotyped virus co-expressing SARS-CoV-2 spike protein and GFP reporter (VSV*ΔG-GFP SARS-CoV-2 S) ^42^. As expected, Huh7 cells overexpressing BTN3A2-L or BTN3A2-S significantly inhibited VSV*ΔG-GFP SARS-CoV-2 S, whereas knockout of BTN3A2 in Huh7 cells (sgBTN3A2-1 Huh7 cells and sgBTN3A2-3 Huh7 cells) enhanced VSV*ΔG-GFP SARS-CoV-2 S replication (Supplementary Figs. S3b and S3c). We performed a syncytia formation assay to test whether BTN3A2 restricts viral entry using Huh7 cells overexpressing BTN3A2-L or BTN3A2-S. We found that overexpression of BTN3A2-L and BTN3A2-S had no apparent effect on Spike protein-mediated syncytia formation (Fig. 3b). These results suggested that BTN3A2 most likely interrupted the attachment step of viral invasion.

**Fig. 3.**
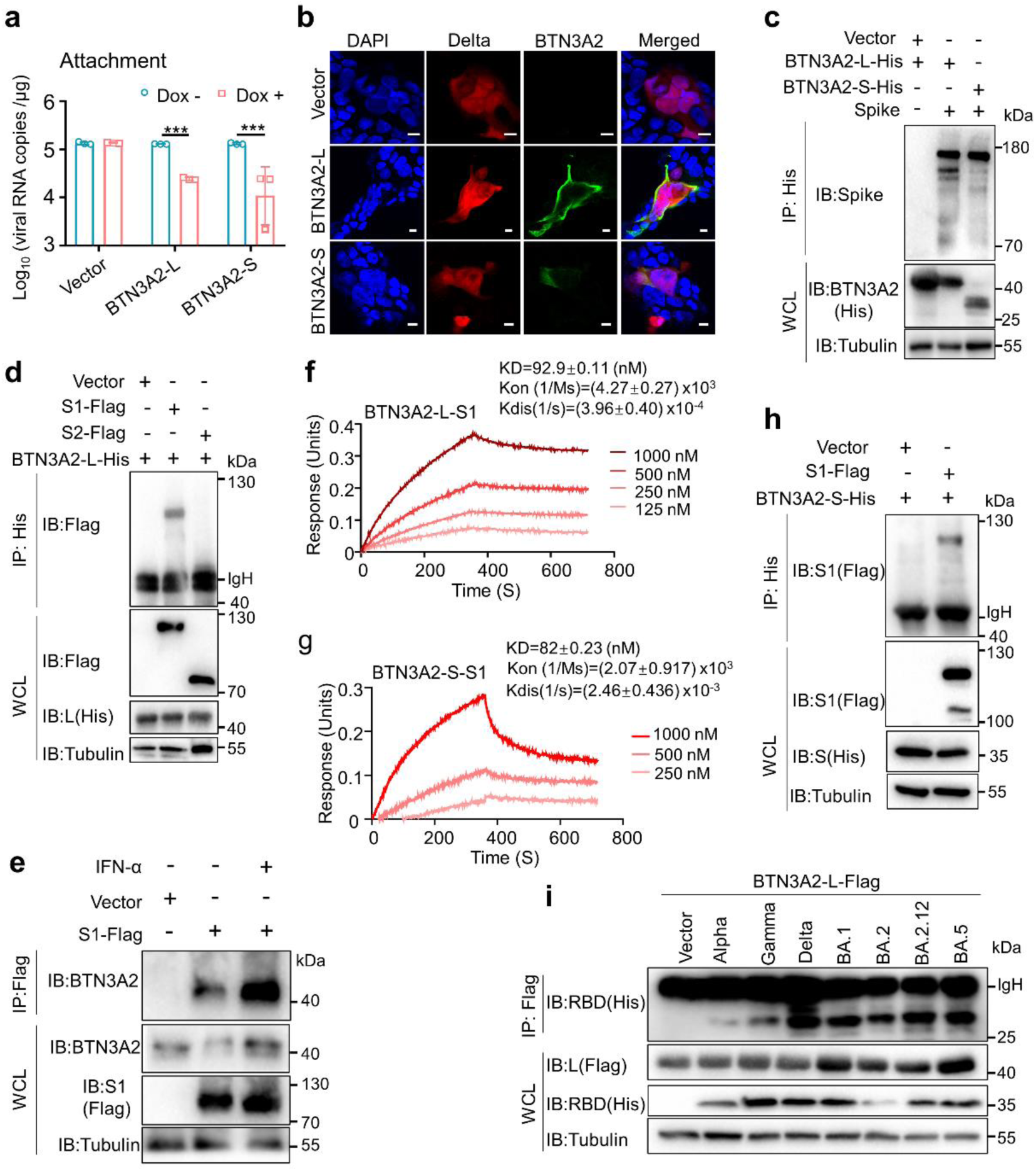
BTN3A2 inhibited SARS-CoV-2 entry and interacted with Spike protein via its RBD. **a**: Stable expression of BTN3A2 suppressed SARS-CoV-2 attachment to the cell. Huh7 cells overexpressing BTN3A2-L (BTN3A2-L Huh7) or BTN3A2-S (BTN3A2-S Huh7) and the control (Vector Huh7) cells were treated with DOX (1 μg/mL) for 24 h, then incubated with SARS-CoV-2 (MOI=1) on ice at 4 °C for 1 h. Cells were washed by PBS three times and harvested to measure viral RNA copies based on SARS-CoV-2 *N* gene level. **b**: Representative immunofluorescence images of cell-cell fusion assay showing no apparent effect of BTN3A2 overexpression (BTN3A2-L and BTN3A2-S) on syncytia formation mediated by Spike of Delta. The HEK293T cells were transfected with Spike of Delta expression vector, and the Huh7 cells were transfected with px-300 for 16 h. The transfected Huh7 and HEK293T cells were co-cultured for 20 h and harvested for immunofluorescence assays. Scale bar, 10 μm. **c**: BTN3A2 interacted with the SARS-CoV-2 Spike protein. HEK293T cells (1×10^7^) were co-transfected with expression vector of BTN3A2 isoform (BTN3A2-L-His or BTN3A2-S-His, each 5 μg) and Spike protein (5 μg) for 48 h. Whole-cell lysates were immunoprecipitated (IP) using anti-His antibody. Immunoblotting (IB) for His (BTN3A2-L-His or BTN3A2-S-His), Spike, and Tubulin was performed using anti-His, anti-Spike, and anti-Tubulin antibodies, respectively. **d**: BTN3A2-L interacted with the S1 subunit but not the S2 subunit of Spike protein. Spike S1 and S2 subunits were Flag-tagged. **e**: Endogenous BTN3A2 interacted with the Spike S1 protein. Huh7 cells (1×10^8^) were transfected with expression vector for Spike S1 protein (S1-Flag, 20 μg) or empty vector (Vector) for 36 h, then cells were treated with or without IFN-α (450 U) for 12 h. Whole-cell lysates were immunoprecipitated (IP) using anti-Flag antibody. Immunoblotting (IB) for BTN3A2, Flag (S1-Flag), and Tubulin was performed using anti-BTN3A2, anti-Flag, and anti-Tubulin antibodies, respectively. **f-g**: Biolayer interferometry analyses of Spike S1 protein binding to immobilized BTN3A2-L-His (**f**) and BTN3A2-S-His (**g**). Different concentrations (1 000, 500, 250, and 125 nM, corresponding to kinetic curves from top to bottom) of Spike S1 protein were used for binding to BTN3A2-L-His and BTN3A2-S-His (lacked 125 nM), respectively. **h**: BTN3A2-S interacted with the Flag-tagged S1 subunit of the Spike protein. **i**: BTN3A2 interacted with Spike RBDs of different SARS-CoV-2 variants. Procedures for IP and IB in (**d**), (**h**) and (**i**) are the same as those in (**c**). Data are representative of three independent experiments with similar results (**b**-**i**). (**a**) Values were presented as mean±SD, and differences between groups were determined by comparing to the Dox-group; ANOVA with Dunnett’s multiple comparisons, ***, *P* < 0.001.

To assess whether BTN3A2 interacts with the SARS-CoV-2 Spike protein, an envelope protein essential for host cell binding and entry^14, 15, 73^, we conducted a co-immunoprecipitation assay. Both BTN3A2 isoforms were co-immunoprecipitated with the Spike protein in HEK293T cells (Fig. 3c). Given the reliance of not only SARS-CoV-2 but also SARS-CoV ^74^ and HCoV-NL63 ^75^ on Spike protein-ACE2 interaction for cell entry, we examined whether BTN3A2 has a similar interaction with the Spike proteins of SARS-CoV and HCoV-NL63. We observed an expected interaction between BTN3A2 and Spike protein of SARS-CoV or HCoV-NL63 (Supplementary Fig. S3d), suggesting a potentially expanded role of BTN3A2 in the related virus infections. Of note, BTN3A2-L interacted with the Spike S1 subunit but not the S2 subunit (Fig. 3d), as further confirmed by the observation that endogenous BTN3A2-L was immunoprecipitated by exogenous Spike S1 overexpression (Fig. 3e) and the biolayer interferometry analysis (KD=92.7±0.111 nM) (Fig. 3f). Similarly, we observed an interaction between BTN3A2-S and S1 subunit (Fig. 3g-h).

Considering a high degree of sequence identity between the extracellular regions of BTN3A2 and its family member (BTN3A1 or BTN3A3), we further tested the potential interaction between BTN3A members with S1. We found that BTN3A1 and BTN3A3 could also interact with S1, but with a lower binding affinity comparing to that of the interaction between BTN3A2 and S1 (Supplementary Fig. S4a-b). We speculated that the lack of the intracellular B30.2 domain of BTN3A2 (relative to BTN3A1 or BTN3A3) might be responsible for its high binding affinity to S1.

The S1 subunit, which contains the RBD, is both an inducer of dominant neutralizing antibodies (nAbs) in the host and a key antigenic target for vaccine development ^41, 76, 77^. Subsequent co-immunoprecipitation assays confirmed that BTN3A2-L interacted with the Spike protein RBD from multiple SARS-CoV-2 variants, including α, γ, δ, BA.1, BA.2, BA.2.12, and BA.5 (Fig. 3i). Similarly, BTN3A2-S could interact with RBD of BA.2 (Supplementary Fig. S3e). These findings established BTN3A2 as a novel cellular binding partner for the SARS-CoV-2 Spike protein specifically at the RBD.

### BTN3A2 competed with ACE2 to bind to RBD

Although several host receptors for SARS-CoV-2 have been identified, only ACE2 interacts with the Spike protein via the RBD ^78^. Thus, we tested whether BTN3A2 isoforms interact with ACE2, thereby modulating ACE2 and Spike protein interactions during viral entry and replication. Both exogenously overexpressed BTN3A2-L (His-tagged) (Fig. 4a-b) and endogenous BTN3A2 could interact with exogenously overexpressed ACE2 according to our co-immunoprecipitation analyses (Fig. 4c). This interaction could be further confirmed by the biolayer interferometry analysis, in which an affinity between BTN3A2-L and ACE2 (KD=92.7±0.359 nM) (Fig. 4d) was observed, indicating a direct association between BTN3A2-L and ACE2. Compared to the interaction between BTN3A2-L and ACE2, a lower affinity was observed between BTN3A2-S and ACE2 (KD=190±0.305 nM) (Fig. 4e) and between the other BTN3A family member (BTN3A1, Supplementary Fig. S4c; BTN3A3, Supplementary Fig. S4d) and ACE2.

**Fig. 4.**
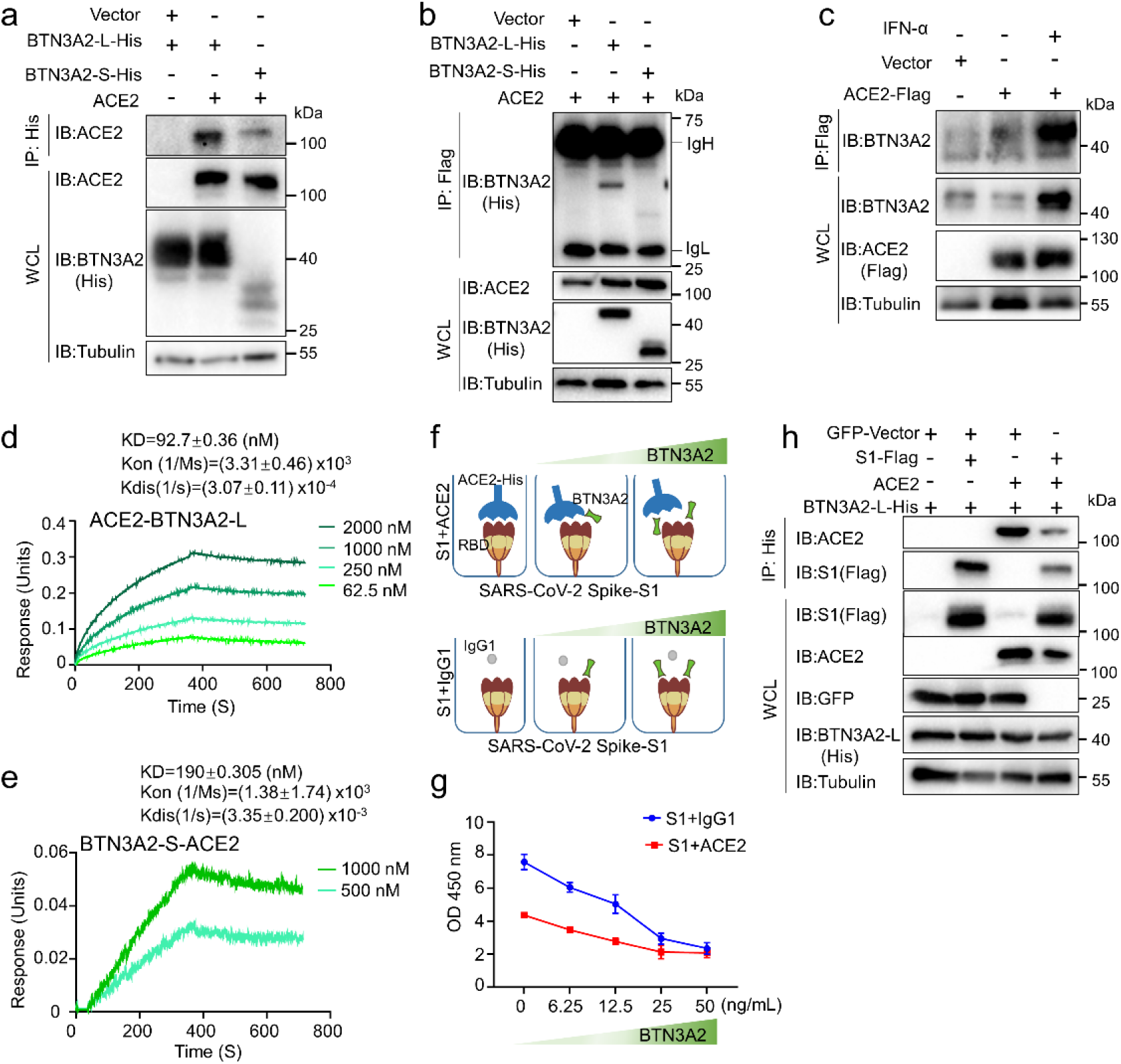
BTN3A2 competed with ACE2 to bind to the Spike protein. **a**: BTN3A2 isoform (BTN3A2-L-His or BTN3A2-S-His) interacted with ACE2. **b**: ACE2 interacted with BTN3A2-L. **c**: Endogenous BTN3A2 interacted with the ACE2. Procedures were similar to Fig. 3e. **d**: Biolayer interferometry analysis of BTN3A2-L binding to immobilized ACE2-Fc. Different concentrations of BTN3A2-L (2 000, 1 000, 250, and 62.5 nM; corresponding to kinetic curves from top to bottom) were used. **e**: Biolayer interferometry analysis of BTN3A2-S binding to immobilized ACE2-Fc. Different concentrations of BTN3A2-S (1 000 and 500 nM; corresponding to kinetic curves from top to bottom) were used. **f-g**: Competition ELISA analysis of competitive binding of BTN3A2-L with Spike RBD (Spike-RBD). **f**: Schematic of competition assays. **g**: Competition between ACE2 (ACE2-His-tagged) and BTN3A2-L (BTN3A2) for immobilized Spike S1 subunit (S1-Fc). ACE2 and S1-Fc protein or human IgG1-Fc and S1-Fc protein were incubated for 12 h, followed by the addition of dilution series of BTN3A2-L-His. Amount of S1 protein remaining in the presence of the competitor was determined by antihuman HRP through a colorimetric readout. **h**: BTN3A2-L competed with ACE2 to bind to the Spike S1. Procedures for IP and IB in (**a-b**) and (**h**) were similar to those in Fig. 3c.

As both ACE2 and BTN3A2 interacted with the Spike protein via its RBD, we further investigated whether BTN3A2 competes with ACE2 for Spike S1 subunit binding. Based on competitive ELISA, the interaction between ACE2 and SARS-CoV-2 Spike was measured in the presence of recombinant BTN3A2-L protein (Fig. 4f). Results showed that Spike-ACE2 binding was greatly diminished under increasing BTN3A2-L concentrations in a dose-dependent manner (Fig. 4g). Co-immunoprecipitation assays demonstrated a decrease in the expression levels of S1and ACE2 under the ectopic expression of BTN3A2-L in HEK293T cells (Fig. 4h), while was in general consistent with a previous report showing a degradation of ACE2 by SARS-CoV-2 Spike protein ^79^ in whole cell lysate (WCL). These findings suggest that BTN3A2 interacts with the Spike through RBD binding, and ACE2 and BTN3A2 compete for binding to the Spike protein.

### BTN3A2 inhibited ACE2 expression

We next explored the potential role of the interactions among BTN3A2, ACE2, and Spike protein in the context of COVID-19 pathology. We first compared the tissue distribution patterns of BTN3A2 and ACE2. Re-analyses of a large-cohort RNA-seq dataset of GTEx human tissues ^59^ showed that *BTN3A2* mRNA expression exhibited a reverse correlation with *ACE2* in the lungs and gastrointestinal tract (small intestine) (Supplementary Fig. S5a), which are major targets of SARS-CoV-2 infection. A similar pattern was also observed in lung tissues and cell lines with or without SARS-CoV-2 infection ^80^ (Supplementary Fig. S5b). We further compared *BTN3A2* co-expression pattern with *ACE2* in different lung cell types using scRNA-seq data from COVID-19 patients ^58^. Our results indicated that, following SARS-CoV-2 infection, *ACE2* mRNA could be detected at a slightly increased level in the lung epithelial cells of COVID-19 patients, but *BTN3A2* expression was substantially up-regulated in the same cell type (Fig. 5a). This observation suggested that the upregulation of *BTN3A2* and *ACE2* mRNA was possibly controlled by different signaling pathways, although BTN3A2 could reduce ACE2 protein expression. Consistently, the SARS-CoV-2-infected lung tissues of NHPs had a lower level of *ACE2* mRNA compared to that of lung tissues from uninfected animals (Supplementary Fig. S5c).

**Fig. 5.**
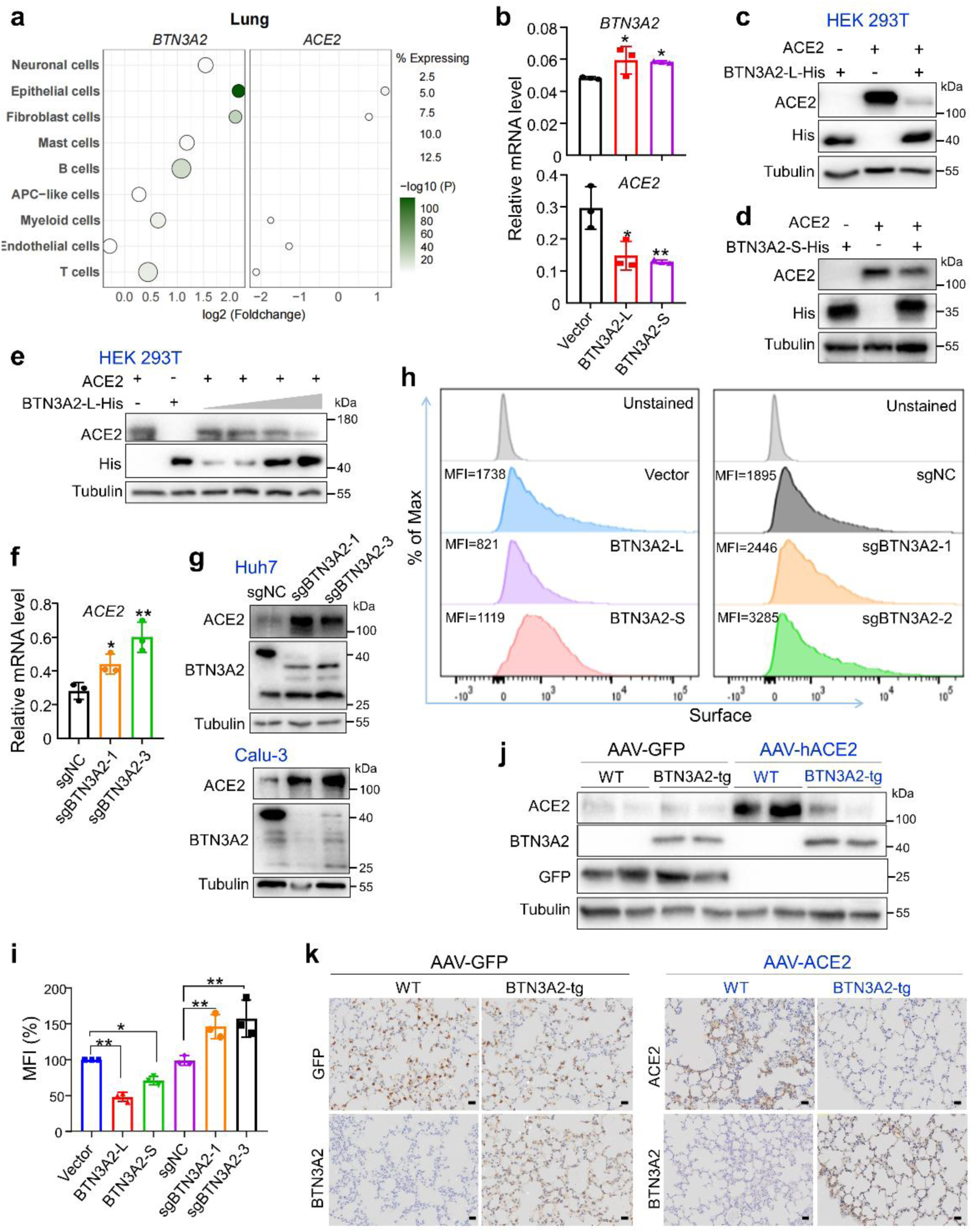
BTN3A2 inhibited ACE2 expression. **a**: Differential expression of *BTN3A2* and *ACE2* in scRNA-seq dataset of lung tissues from COVID-19 patients and uninfected individuals. Original dataset was reported in Melms et al. ^58^. Significance was assessed by two-sided Wilcox rank sum test. **b**: Stable expression of BTN3A2 decreased *ACE2* mRNA level in Huh7 cells. RNA was extracted from Huh7 cells overexpressing BTN3A2-L and BTN3A2-S, or control (Vector Huh7) cells (each group 1×10^5^ cells). *BTN3A2* and *ACE2* mRNA levels were measured by qRT-PCR, with normalization to *GAPDH*. **c-d**: Overexpression of BTN3A2-L (**c**) and BTN3A2-S (**d**) suppressed ACE2 protein expression in HEK293T cells. Cells (5×10^5^) were transfected with indicated expression vector or empty vector (each 1.25 μg) for 48 h, then harvested for western blotting. **e**: Dose-dependent inhibitory effect of BTN3A2-L on ACE2 protein expression. HEK293T cells (5×10^5^) were transfected with indicated expression vector or empty vector (each 0.5 μg), together with increased BTN3A2-L-His (0.25, 0.50, 0.75, and 1.50 μg, with empty vector to reach a total amount of 2.5 μg of plasmid) for 48 h, then harvested for western blotting. **f**: Knockout of BTN3A2 increased *ACE2* mRNA level in Huh7 cells. The procedure was similar to that in (b). **g**: Knockout of BTN3A2 promoted ACE2 protein expression in Huh7 (*upper*) and Calu-3 (*lower*) cells. **h-i**: Flow cytometry was conducted to show surface ACE2 in Calu-3 cells with BTN3A2 overexpression (BTN3A2-L and BTN3A2-S) or knockout (sgBTN3A2-1 and sgBTN3A2-2), together with the control cells (vector and sgNC) and unstained cells. Cell surface expression of ACE2 was measured (**h**), and quantification of mean fluorescence intensity (MFI (%), *n*=3) was based on three independent experiments (**i**). **j**: ACE2 in BTN3A2-tg and WT mice infected with indicated AAV-GFP and AAV-hACE2. Immunoblotting for ACE2, BTN3A2, GFP, and Tubulin was performed using anti-ACE2, anti-BTN3A2, anti-GFP, and anti-Tubulin antibodies, respectively. **k**: Representative images of immunohistochemical staining of ACE2 and BTN3A2 in lung tissues in (**j**). Scale bar, 100 μm. Data were representative of three independent experiments with similar results (**b-k**). Values were presented as mean±SD (**b**, **f**, and **i**). Significance was determined compared to Vector (**b** and **i**) or sgNC (**f** and **i**). ns, not significant; *, *P* <0.05; **, *P*<0.01; ANOVA with Dunnett’s multiple comparisons.

To further determine whether BTN3A2 affects ACE2 expression, we assessed whether stable expression of *BTN3A2* in Huh7 cells altered the expression of *ACE2* mRNA. Our results indicated that *ACE2* mRNA expression was significantly decreased in Huh7 cells with stable expression of BTN3A2-L (BTN3A2-L Huh7 cells) and BTN3A2-S (BTN3A2-S Huh7 cells) (Fig. 5b). Consistently, total ACE2 protein expression was markedly decreased in HEK293T cells overexpressing both BTN3A2-L and ACE2 compared to cells overexpressing ACE2 only (Fig. 5c). However, BTN3A2-S had minor effect on inhibiting ACE2 expression in HEK293T cells (Fig. 5d) compared to BTN3A2-L. The inhibitory effects of BTN3A2-L on ACE2 expression were dose-dependent (Fig. 5e). Knockout of BTN3A2 in Huh7 cells (sgBTN3A2-1 Huh7 cells and sgBTN3A2-3 Huh7 cells) led to a significant increase in both *ACE2* mRNA (Fig. 5f) and protein levels (Fig. 5g). Using flow cytometry, we further assessed whether BTN3A2 alters ACE2 expression on the cell surface. Notably, ACE2 expression on the cell surface was significantly decreased in Calu-3 cells with stable expression of BTN3A2-L (BTN3A2-L Calu-3 cells) and BTN3A2-S (BTN3A2-S Calu-3 cells), while knockout of BTN3A2 in Calu-3 cells (sgBTN3A2-1 Calu-3 cells and sgBTN3A2-3 Calu-3 cells) increased the surface level of ACE2 (Fig. 5h-i). Similar results were observed in genetic modified Huh7 cell lines (Supplementary Fig. S5d-e), indicating that BTN3A2 inhibits ACE2 expression *in vitro*.

As *BTN3A2* is a primate-specific gene ^26^, we used BTN3A2-tg mice to further characterize the level of ACE2 *in vivo*. WT C57BL/6J mice were used as a control. Mice were intranasally infected with AAV-hACE2 (AAV carrying the human *ACE2* gene, AAV-hACE2 group) or AAV-EGFP (AAV-EGFP group, control group). Immunoblot analysis showed that EGFP levels were similar between the BTN3A2-tg and WT C57BL/6J mice (WT control) (Fig. 5j), indicating effective viral vector-mediated gene expression. Consistent with the cellular assays and inhibitory effects of BTN3A2 on ACE2 expression, our results further showed that the ACE2 protein level was significantly lower in BTN3A2-tg mice relative to WT C57BL/6J mice (Fig. 5j). The AAV vectors achieved efficient EGFP and ACE2 expression in the lung tissues of WT mice at 14 dpi with AAV (Fig. 5k and Supplementary Fig. S5f). Conversely, ACE2 was barely detected in the lung tissues of BTN3A2-tg mice (Fig. 5k and Supplementary Fig. S5f). Collectively, these results suggest that BTN3A2 suppression of ACE2 also occurs *in vivo*.

### BTN3A2 suppressed SARS-CoV-2 infection by reducing ACE2

We further evaluated whether the suppression of ACE2 expression by BTN3A2 is responsible for BTN3A2-mediated SARS-CoV-2 restriction. We intranasally injected AAV9-hACE2 in WT C57BL/6J and BTN3A2 transgenic (BTN3A2-tg) heterozygous mice and observed for two weeks, followed by SARS-CoV-2 challenge and sacrifice for tissue collection at 3 dpi (Fig. 6a). Quantitative analysis of viral loads, specifically targeting the nucleoprotein gene (*N*) and envelope gene (*E*), revealed a 3-fold to 6-fold decrease in lung tissue viral concentrations in BTN3A2-tg mice compared to their WT counterparts (Fig. 6b). Of the seven infected WT C57BL/6J mice examined, five exhibited detectable levels of viral subgenomic RNA in their lung tissues, including the subgenomic E (sgE) gene, a marker of infectious virus ^38, 81^. In contrast, none of the six BTN3A2-tg mice showed detectable levels of sgE gene. Further immunohistochemical staining of the SARS-CoV-2 nucleoprotein (N) showed a consistent reduction in viral protein levels in the lung tissues of BTN3A2-tg mice compared to the infected WT C57BL/6J mice (Fig. 6c). Moreover, histological analysis showed that the lung tissues of the SARS-CoV-2 infected BTN3A2-tg mice had milder damage, including the widening, fusion, and increased solidity of the pulmonary septa, compared to the infected WT C57BL/6J mice (Fig. 6d), suggesting a protective role of BTN3A2 in counteracting SARS-CoV-2 infection. Similarly, we observed consistent and effective expression of hACE2 in the lung tissues of the WT group, while the protein level of ACE2 was greatly reduced in the lung tissues of the BTN3A2-tg mice, as expected (Fig. 6e). This pattern was further validated by immunohistochemical analysis of lung tissues from both groups: ACE2 was consistently and highly expressed in the upper and lower respiratory tract of WT mice (Fig. 6f) but was barely detected in the respiratory tract of BTN3A2-tg mice. Collectively, these results suggest that BTN3A2 can restrict SARS-CoV-2 by suppressing ACE2-mediated viral entry *in vivo*.

**Fig. 6.**
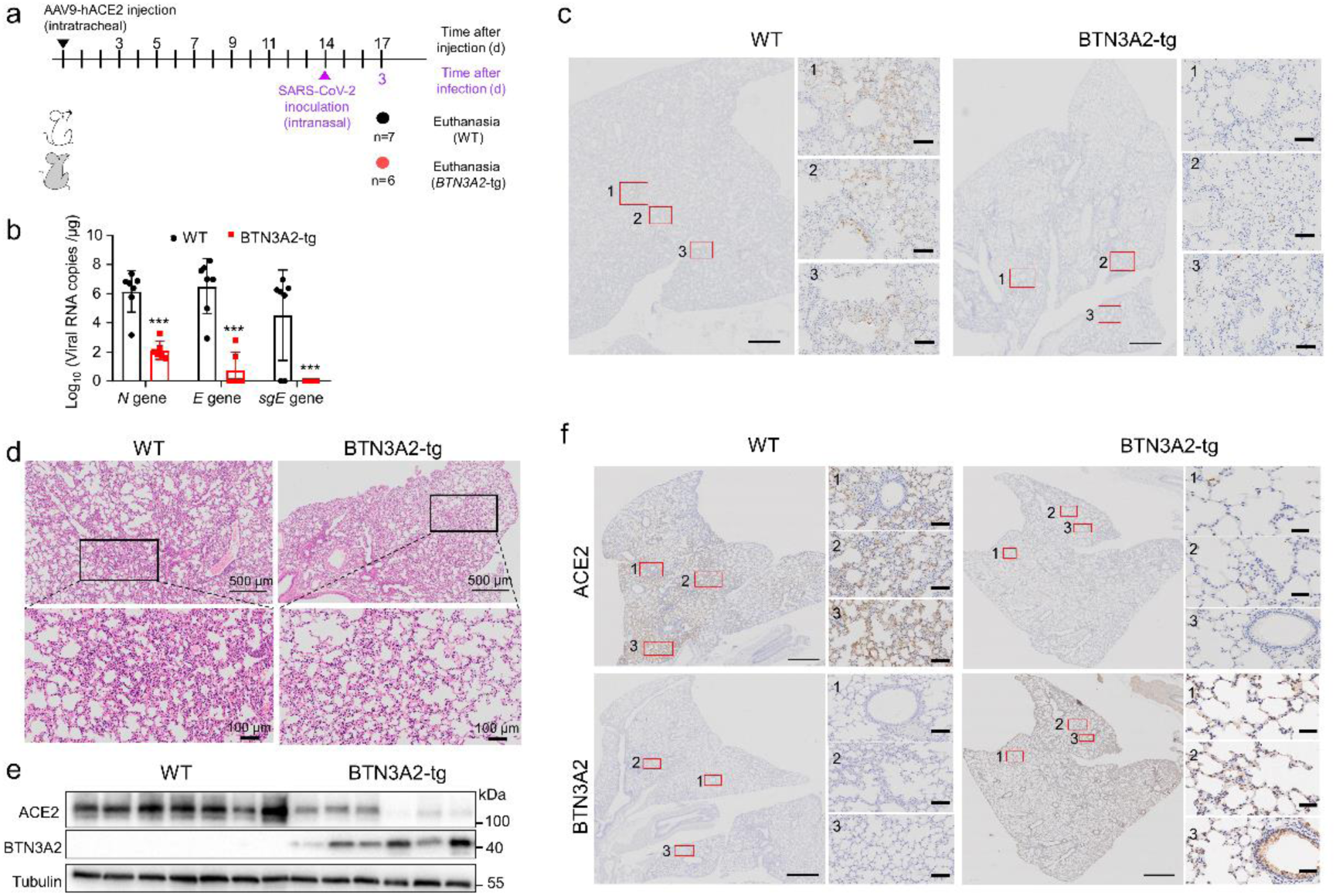
BTN3A2 suppressed ACE-mediated SARS-CoV-2 entry. **a**: Schematic of animal experiments with SARS-CoV-2 inoculation. BTN3A2-tg (*n*=6) and WT mice (C57BL/6J, *n*=7) were intranasally infected with AAV9-hACE2 (2×10^12^ viral genome copies) for 14 days, with all animals then infected with BA.2 (5×10^4^ TCID_50_). All mice were euthanized at 3 dpi for tissue collection. **b**: Viral RNA copies in lung tissues of SARS-CoV-2-infected mice at 3 dpi. Each dot indicates log copies of viral genomic N gene, E gene, and subgenomic E (sgE) of a mixed sample of five lung lobes from an individual mouse. Viral copies are presented as mean±SD. ***, *P*<0.001; ANOVA with Dunnett’s multiple comparisons. **c**: Representative image of immunohistochemical staining of SARS-CoV-2 N protein in lung tissues of WT and BTN3A2-tg mice at 3 dpi. Scale bars: 500 μm for entire lung lobe section (*left*) and 100 μm for enlarged view of boxed areas labeled by numbers in entire section (*right*). **d**: Representative of image of H&E staining of lung sections in SARS-CoV-2-infected WT and BTN3A2-tg mice. **e**: AAV-mediated hACE2 expression in lungs tissues of WT (C57BL/6J, *n*=7) and BTN3A2-tg mice (*n*=6) at 3 dpi. **f**: Representative image of immunohistochemical staining for ACE2 and BTN3A2 in lung tissues of WT and BTN3A2-tg mice at 3 dpi. Scale bars were same as those in (**c**).

## Discussion

Understanding the pathobiology of SARS-CoV-2 infection is essential for developing effective treatments. Various studies have examined the host immune response in relation to SARS-CoV-2 infection ^50, 82, 83^. Specifically, the balanced regulation of interferon signaling and immune response has been shown to substantially influence disease severity and outcomes of COVID-19 ^6, 84^. In the current study, we identified BTN3A2 as a novel factor involved in the defense against SARS-CoV-2 infection. Notably, BTN3A2 was up-regulated upon SARS-CoV-2 infection and directly interacted with the RBD of the Spike protein. This interaction was confirmed through both cell-free binding assays utilizing recombinant proteins and co-immunoprecipitation assays conducted in cellular environments. This led to competitive interference with the interaction between ACE2 and the RBD of the Spike protein, as reported previously ^13, 85^. Given that BTN3A2 is a single transmembrane protein localized to the cellular membrane ^86^, it is plausible that a complex between BTN3A2 and the Spike protein may effectively obstruct the binding of the Spike protein to ACE2 on the cellular surface. Remarkably, BTN3A2-S, which lacks a transmembrane domain, demonstrated greater efficacy than BTN3A2-L in inhibiting the attachment of the Spike protein to the cellular surface, thereby reducing SARS-CoV-2 infection. We hypothesize that secreted BTN3A2-S ^26^ may possess enhanced inhibitory effects against SARS-CoV-2 Spike binding to ACE2, akin to the function of neutralizing antibodies. Investigating the structural characteristics and molecular dynamics of the BTN3A2-Spike protein complex could offer substantial insights for the engineering of short peptides that retain the full competitive capacity of BTN3A2 in challenging ACE2 for Spike protein binding.

Reducing ACE2 expression has been considered a potentially efficacious host-directed prophylaxis for protection against SARS-CoV-2 infection ^23, 83^. Beyond the competitive effects of BTN3A2 on ACE2-Spike binding, our study demonstrated that BTN3A2 interacted with ACE2, leading to a further decrease in ACE2 expression. Overexpression of BTN3A2 led to a reduction in *ACE2* mRNA and protein levels, which further inhibited ACE2-mediated SARS-CoV-2 infection, both *in vitro* and *in vivo*. While the exact mechanism by which BTN3A2 regulates ACE2 expression remains to be determined, the apparent dual role of BTN3A2 in regulating ACE2 expression and function is consistent with the concept that targeting universal host factors essential for viral replication offers advantages in terms of broad-spectrum effectiveness and reduced drug resistance ^87, 88^.

The ongoing COVID-19 pandemic, coupled with the emergence of new SARS-CoV-2 variants, necessitates the development of novel broad-spectrum vaccines and antiviral therapies. Recent studies have identified several potential targets for SARS-CoV-2 intervention, including the transcriptional regulator BRD2 ^83^, aryl hydrocarbon receptor (AhR) ^8^, and host cysteine-aspartic protease caspase-6 ^89^, for which specific compounds have been characterized. Given the observed protective effects of BTN3A2 against SARS-CoV-2 infection, we speculate that BTN3A2 may be a viable host factor candidate with broad antiviral potential. Indeed, pre-incubation of soluble BTN3A2 protein with host cells resulted in decreased viral entry. Notably, interactions between BTN3A2 and the Spike protein RBD of various SARS-CoV-2 variants of concern (VOCs) (e.g., prototype, α, γ, δ, and omicron) remained largely stable despite the rich diversity and evolution of viral sequences. These findings suggest the anti-coronavirus activity of BTN3A2 may be both conserved and efficacious across multiple SARS-CoV-2 VOCs.

The primate-specific *BTN3A2* gene ^26^ is highly expressed in the brain. Thus, it may have a role in the development of long COVID and brain fog, conditions characterized by a multitude of persistent neurological symptoms that last for months after SARS-CoV-2 infection ^90^. Currently, the pathophysiological mechanisms associated with neurological symptoms derived from SARS-CoV-2 infection remain to be elucidated ^91^. We recently showed that the S2 subunit of the SARS-CoV-2 Spike protein modulates γ-secretase and enhances Aβ production, which may contribute to the neurological alterations observed in COVID-19 patients ^45^. Respiratory infection with SARS-CoV-2 can elicit neuroinflammation and dysfunction in neural cells and myelin ^92^. Modulation of the immune environment within the central nervous system, particularly in scenarios where neuroinflammation is triggered by inflammation in the respiratory system, may influence the neuroprotective and neuroplastic functions of specific immune cell populations, including γδ T cells ^93^, following COVID-19 ^91^. Therefore, given the broad immune cell expression of BTN3A2 and its involvement in γδ T cell regulation ^27, 94^, determining the roles of BTN3A2 in T cell-mediated immune responses, neuroinflammation, and long COVID-19 would be worthwhile.

This study has several limitations. Firstly, although we presented multiple lines of evidence to show the active role of BTN3A2 as a key host factor in combating SARS-CoV-2 infection, how BTN3A2 was upregulated upon SARS-CoV-2 infection, and which signaling pathways were activated in this process remain unknown. It could be possible that induction of the interferon system by SARS-CoV-2 led to an increased expression of *BTN3A2*, as shown by the fact that the *BTN3A2* expression was upregulated in IFN-α treated Calu-3 cells (Supplementary Fig. S1f) and Huh7 cells (Fig. 3e and Fig. 4c). Focused study should be performed to refine this mechanism. Secondly, we showed evidence that BTN3A2 interacts with the Spike proteins of SARS-CoV-2 VOCs, indicating a potential broad-spectrum antiviral role of BTN3A2. This speculation deserves further validations. Thirdly, as a single-pass transmembrane protein on the host cell membrane, whether BTN3A2, by competitively binding to the RBD of the Spike protein, could function as an alternative receptor for SARS-CoV-2 and assist in viral invasion, is an arising question from this study.

In conclusion, we identified BTN3A2 as a novel host factor with protective effects against SARS-CoV-2 infection. Thus, BTN3A2 holds considerable potential as a therapeutic drug for mitigating the impact of SARS-CoV-2 VOCs. However, subsequent research is warranted to explore its potential role in the occurrence of neurological symptoms, commonly referred to as “brain fog”, in COVID-19 patients.

## Supporting information

Table S1 and SOM 20240724

## Contributors

L.X., N.S., Y.-G.Y. conceptualized the study. L.X., D.Y., M.X., Y.L., L.-X.Y., Q-C.Z., X.-L.F., M.-H.L. performed the experiments. D.Y., L.-X.Y., Q-C.Z. verified the underlying data, L.X., D.Y. and M.X. analyzed the data. L.X., D.Y., M.X., Y.-G.Y. wrote the manuscript. Y.-G.Y. and N.S. supervised the study and revised the manuscript. All authors have given approval to the final version of the manuscript.

## Data Sharing Statement

Data reported in this paper will be shared by the lead contact upon reasonable request. This paper does not report original code.

## Acknowledgments

We thank Drs. Yu-Lin Yao and Yong Wu for technical discussions. We are grateful to Profs. Qihui Wang and Yong-Tang Zheng for sharing the VSV-G pseudotyped viruses.

