## Supplementary material for "Primate-specific BTN3A2 protects against SARS-CoV-2 infection by interacting with and reducing ACE2": Table S1 and SOM 20240724

**Supplementary Table S1 Primers and vectors used in this study**

| Primer | Sequence (5'-3') ^a^ | Restriction endonuclease | Application and vector |
| --- | --- | --- | --- |
| *For humans* | |  |  |
| sgBTN3A2-1F | CACCGATACAGTGGAGCAACGCCAA |  | PCR for constructing sgRNA vector (sgBTN3A2-1) using lentiCRISPRv2 vector |
| sgBTN3A2-1R | AAACTTGGCGTTGCTCCACTGTATC |  |  |
| sgBTN3A2-3F | CACCGGATCATGAGAGGCGGCTCCG |  | PCR for constructing sgRNA vector (sgBTN3A2-3) using lentiCRISPRv2 vector |
| sgBTN3A2-3R | AAACCGGAGCCGCCTCTCATGATCC |  |  |
| BTN3A2-L-F | TGCCTATCTTGCTGCTGCTT |  | Analytical quantitative real-time PCR (qRT-PCR) |
| BTN3A2-L-R | AGGCTTATTTCCCGCTCTGT |  |  |
| BTN3A2-S-F | TGGTTGCAGATGGAGTGGGC |  | Analytical qRT-PCR |
| BTN3A2-S-R | CTGGAGGCTCTCTGCGATGG |  |  |
| ACE2-F | GAGGATGTGCGAGTGGCTA |  | Analytical qRT-PCR |
| ACE2-R | ATGGCCTTTTCAACTTCAG |  |  |
| GAPDH-F | CAACTACATGGTTTACATGTTC |  | Analytical qRT-PCR |
| GAPDH-R | GCCAGTGGACTCCACGAC |  |  |
| BTN3A2_F-HA | G***GAATTC***GGATGAAAATGGCAAGTTCCCTGGC | *Eco*RI | PCR for constructing BTN3A2-HA expression vector (BTN3A2-L-HA and BTN3A2-S-HA vector) using pCMV-HA vector |
| BTN3A2_R-HA | CCG***CTCGAG***TCAGGCTGACTTATTGGTATCG | *Xho*I |  |
| BTN3A2_F-pLVX | ATTT***GCGGCCGC***ATCGCCTGGAGAAGGATCCGCG | *Not*I | PCR for constructing pLVX-BTN3A2 expression vector (pLVX-BTN3A2-L and pLVX-BTN3A2-S vector) using pLVX-Tight-Puro vector |
| BTN3A2_R-pLVX | G***GAATTC***GGCTGACTTATTGGTATCGGACG | *Hin*dIII |  |
| *For monkeys* | | | |
| maBTN3A2-L-F | TGAGCAAGAGATGAAAGAACGG |  | Analytical qRT-PCR |
| maBTN3A2-L-R | TTTCTCCTCTTGAGTTCCTCCTG |  |  |
| maGAPDH-F | CTCCTGTTCGAGAGTCAGCC |  | Analytical qRT-PCR |
| maGAPDH-R | GCCCAATACGACCAAATCCG |  |  |
| maACE2-F | TCCTTACCAGTCCCCCGTTA |  | Analytical qRT-PCR |
| maACE2-R | GACGACAATGCCAGCCACTA |  |  |

^a^ Restriction endonuclease sites introduced by PCR are underlined and italicized.

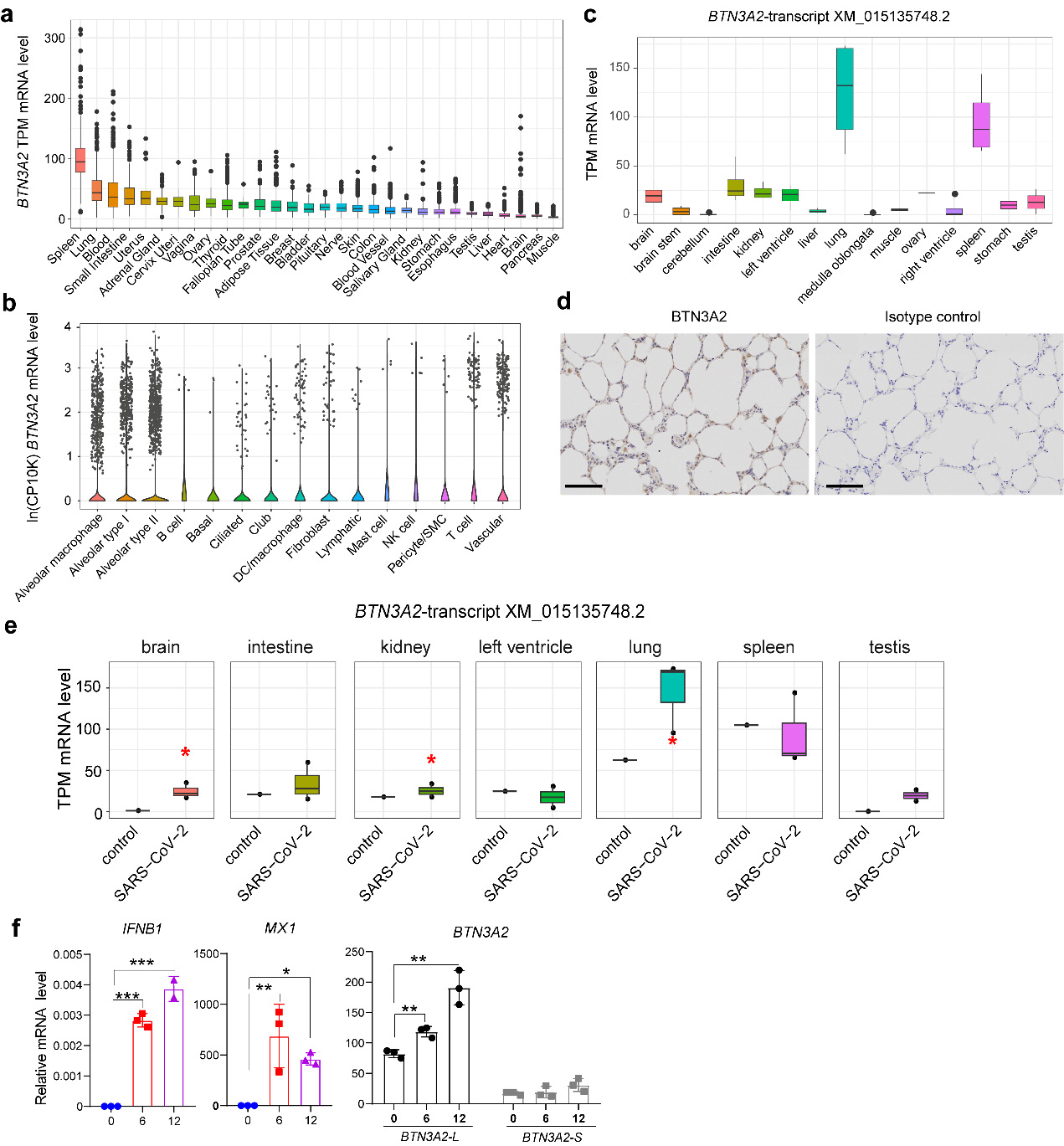

**Supplementary Fig. S1 Expression of *BTN3A2* in human tissues based on GTEx dataset and in Calu-3 cells treated with or without IFN-α.**

**a**: Dot plot showing *BTN3A2* mRNA levels in different human tissues.

**b**: *BTN3A2* mRNA levels in different cell types measured by single-cell transcriptome of healthy human lung tissues. Original data in (a) and (b) were obtained from the GTEx Portal ([www.gtexportal.org](http://www.gtexportal.org)) ^1, 2^. TPM: transcripts per million. CP10K: counts per 10k+ 1.

**c**: *BTN3A2* mRNA levels in 15 tissues from rhesus macaques. Original dataset was reported in Gao et al. ^3^.

**d**: Representative images of immunohistochemical analysis of rhesus macaque lung sections. Cells expressing BTN3A2 (in brown) include epithelial cells and pneumocytes. Scale bars, 100 μm.

**e**: *BTN3A2* mRNA levels in seven tissues from rhesus macaques infected with SARS-CoV-2 (*n*=3) and healthy controls (*n*=1). Original dataset was reported in Gao et al. ^3^. Significance was assessed by two-sided *t*-test; *, *P* < 0.05.

**f**: Upregulation of *BTN3A2* mRNA levels in Calu-3 cells treated with IFN-α. Calu-3 cells were treated with IFN-α (45 U) for indicated times before harvest for total RNA isolation. The mRNA levels of *IFNB1*, *MX1* and *BTN3A2* were measured by qRT-PCR, with normalization to *GAPDH*. Significance was determined by comparing to cells at 0 h for each group. **, *P* < 0.01; ***, *P* < 0.001; ANOVA with Dunnett’s multiple comparisons.

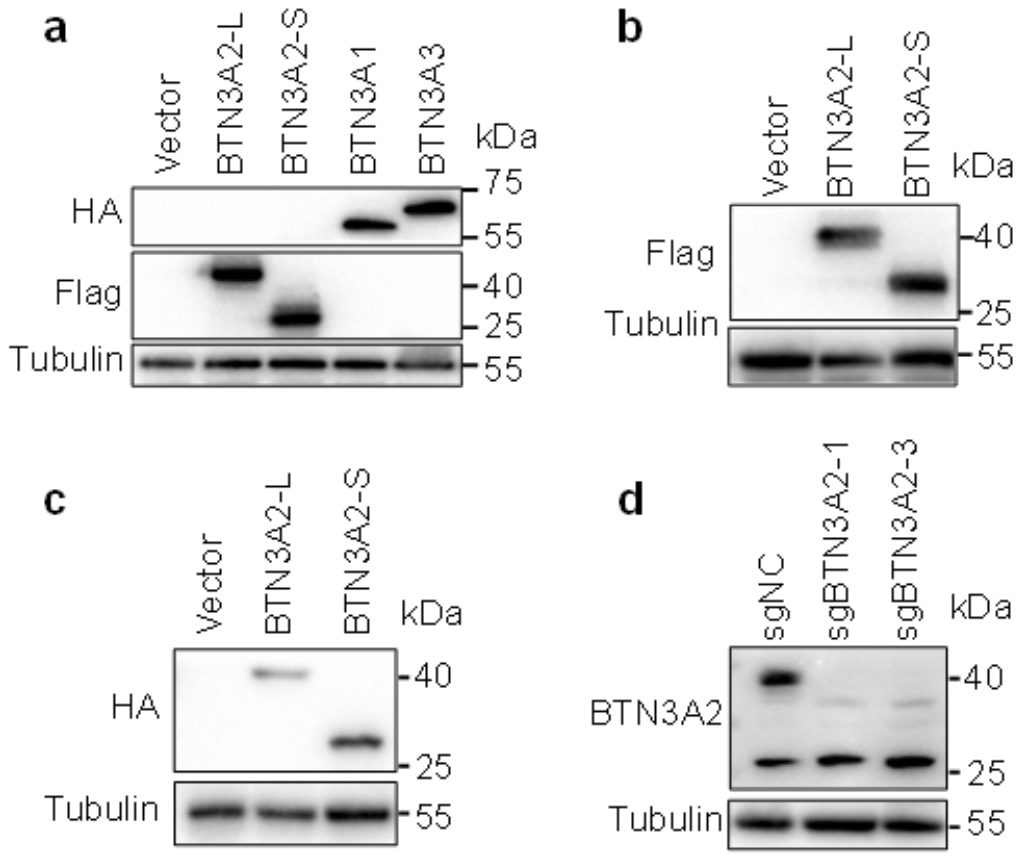

**Supplementary Fig. S2 Successful overexpression or knockout of BTN3A2 in different cells.**

**a**: Successful overexpression BTN3A in ACE2-A549 cells. ACE2-A549 cells (5×10^5^) were transfected with indicated expression vector or empty vector (each 2.5 μg) for 24 h before harvest. Cell lysates were analyzed by western blotting. BTN3A2-L-Flag, BTN3A2-S-Flag, BTN3A1-HA, BTN3A3-HA, and Tubulin were detected using anti-Flag, anti-HA, and anti-Tubulin antibodies, respectively.

**b**: Successful overexpression of BTN3A2-Flag in ACE2-HEK293T cells.

**c**: Successful expression of BTN3A2 isoforms in Huh7 cell line. Cells (5×10^5^) were treated with DOX (1 μg/mL) for 24 h before harvest.

**d**: Successful knockout of BTN3A2 in Huh7 cells using siRNAs. Indicated cells (5×10^5^) were cultured for 24 h before harvest.

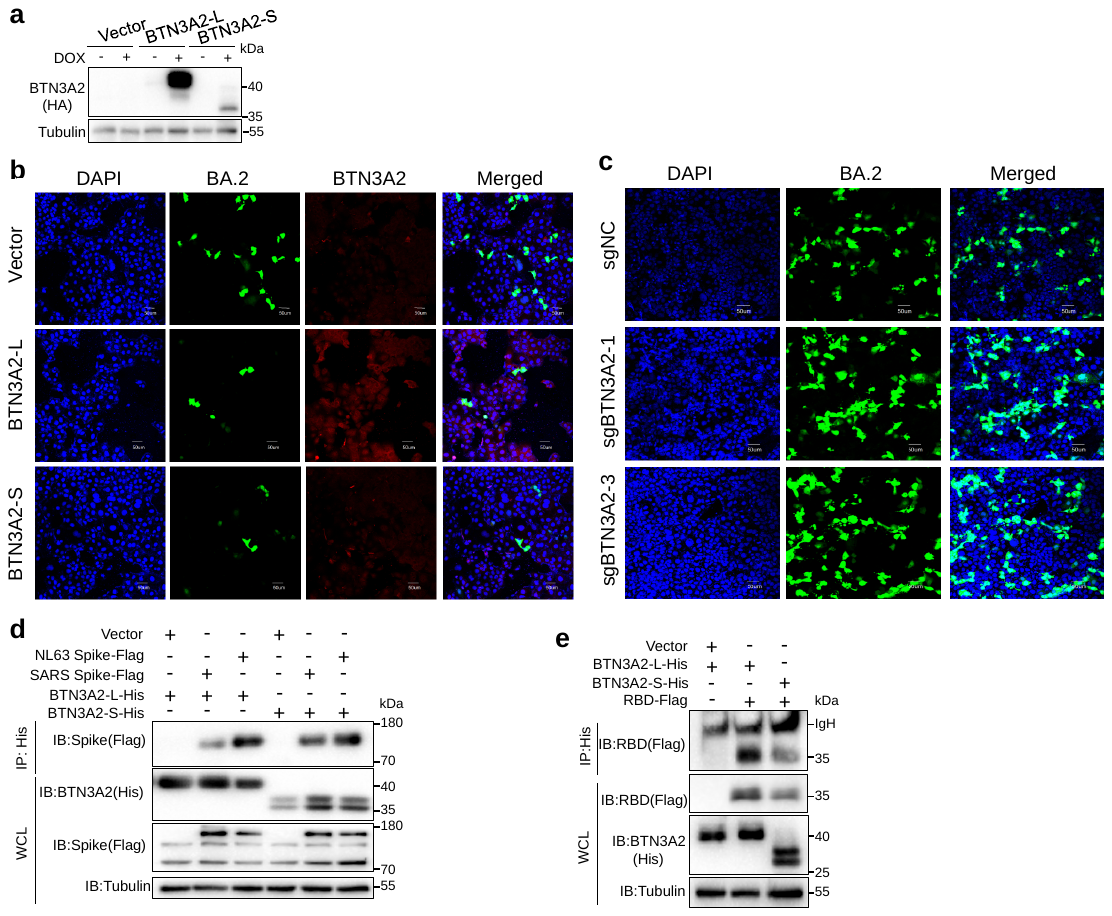

**Supplementary Fig. S3** **BTN3A could interact with Spike.**

**a**: The indicated Huh 7 cell lines (each 5×10^5^ cells) were treated with or without DOX (1 μg/mL) for 24 h before harvest.

**b**: Representative images of immunofluorescence showing a restricted replication of VSV*ΔG-GFP SARS-CoV-2 S in Huh7 cells with BTN3A2 overexpression. Huh7 cells with BTN3A2-L overexpression (BTN3A2-L Huh7 cells), BTN3A2-S overexpression (BTN3A2-S Huh 7 cells), and the control (Vector Huh 7) cells were treated with DOX (1 μg/mL) for 24 h, then were infected with VSV*ΔG-GFP SARS-CoV-2 S for another 24 h before harvest for immunofluorescence assay. VSV*ΔG-GFP SARS-CoV-2 S, a VSV-G pseudotyped virus co-expressing SARS-CoV-2 spike protein BA.2 and GFP reporter.

**c**: Representative images of immunofluorescence showing an increased replication of VSV*ΔG-GFP SARS-CoV-2 S in Huh7 cells with BTN3A2 knockout. sgBTN3A2-1 Huh7 cells, sgBTN3A2-3 Huh7 cells and the control cells (sgNC Huh7) were infected with VSV*ΔG-GFP SARS-CoV-2 S for 24 h before harvest for immunofluorescence assay. Cells were fixed and nuclei were counterstained with DAPI. Scale bar: 50 μm.

**d**: BTN3A2 interacted with the Spike proteins of SARS and NL63.

**e**: BTN3A2-L and BTN3A2-S interacted with BA.2 RBD protein.

Procedures for IP and IB in (**d**) and (**e**) are the same as those in Fig. 3c.

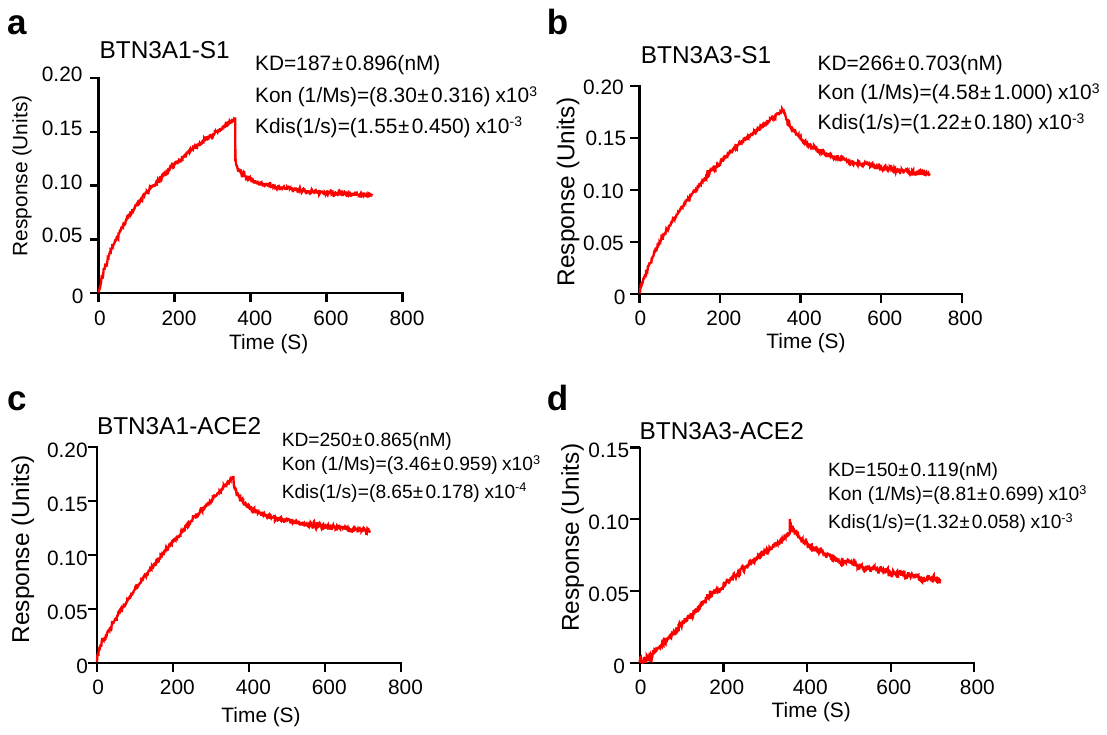

**Supplementary Fig. S4 BTN3A could interact with ACE2 and Spike S1.**

**a-b**: Biolayer interferometry analyses of Spike S1 protein binding to immobilized BTN3A1-His (**a**) and BTN3A3-His (**b**).

**c-d**: Biolayer interferometry analyses of ACE2 protein binding to immobilized BTN3A1-His (**c**) and BTN3A3-His (**d**).

1 000 nM of Spike S1 protein was used in each assay.

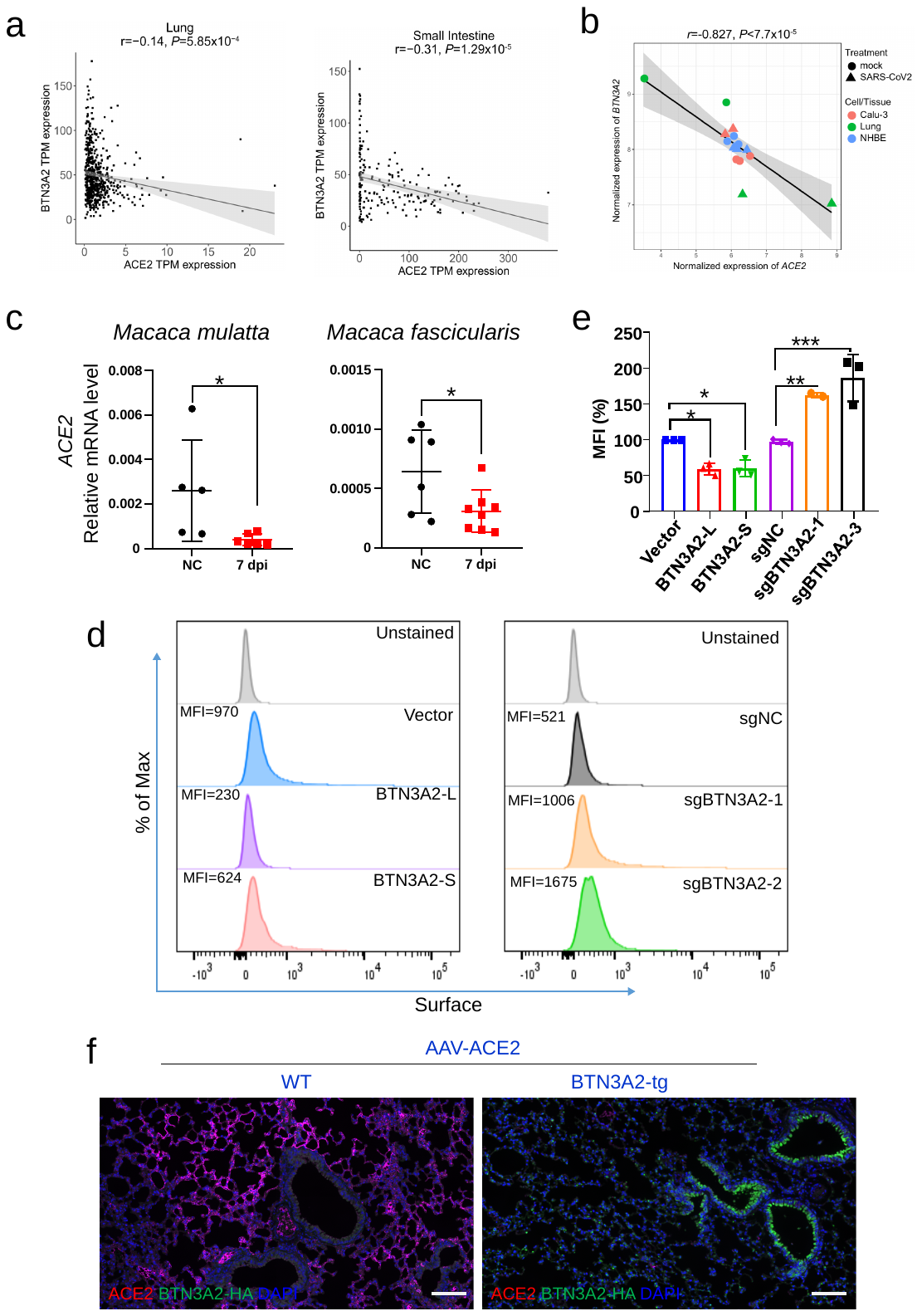

**Supplementary Fig. S5 BTN3A2 affected ACE2 expression.**

**a**: The *BTN3A2* mRNA level was reversely correlated with *ACE2* mRNA level in lung (*left*) and small intestine (*right*) tissues. Each point represents mRNA abundance of a subject from GTEx ^1^.

**b**: The *ACE2* mRNA level was reversely correlated with *BTN3A2* mRNA level in Calu-3 cells, primary human lung epithelial (NHBE) cells, and lung tissues with or without SARS-CoV-2 infection. Original dataset was reported in Sun et al. ^4^. Linear regression line is in black, and 95% compatibility interval is shaded in gray.

**c**: The *ACE2* mRNA levels were down-regulated in lung tissues of rhesus (*left*) and cynomolgus monkeys (*right*) challenged with or without SARS-CoV-2 for 7 days. *ACE2* mRNA level was measured by qRT-PCR and normalized to *GAPDH*. dpi, day post infection.

**d-e**: Flow cytometry analyses of surface ACE2 in Huh7 cells with BTN3A2 overexpression (BTN3A2-L and BTN3A2-S) or knockout (sgBTN3A2-1 and sgBTN3A2-2), together with the control cells (vector and sgNC) and unstained cells. See Fig. 5h-i.

**f**: Representative images of immunofluorescence staining of ACE2 and BTN3A2 in lung tissues in Fig. 5j. Double-immunostaining of lung slices with anti-ACE2 (red) and anti-HA (green) antibodies to indicate expression of ACE2 and BTN3A2-HA, respectively. Scale bar, 100 μm.

Expressional correlation between *BTN3A2* and *ACE2* in (**a-b**) was calculated using Pearson’s correlation analysis. Significance between groups in (**c**) was determined by two-tailed Student’s unpaired *t* test. (**e**) Data were presented as mean±SD of three independent experiments, and significance was determined by ANOVA with Dunnett’s multiple comparisons. *, *P* < 0.05; **, *P* < 0.01; ***, *P* < 0.001.
